# Baicalein suppresses Tau fibrillization by sequestering oligomers

**DOI:** 10.1101/704536

**Authors:** Shweta Kishor Sonawane, Abhishek Ankur Balmik, Debjyoti Boral, Sureshkumar Ramasamy, Subashchandrabose Chinnathambi

## Abstract

Alzheimer’s disease (AD) is a neurodegenerative disorder caused by protein misfolding, aggregation and accumulation in the brain. A large number of molecules are being screened against these pathogenic proteins but the focus for therapeutics is shifting towards the natural compounds as aggregation inhibitors, mainly due to their minimum adverse effects. Baicalein is a natural compound belonging to the class of flavonoids isolated from the Chinese herb *Scutellaria baicalensis*. Here we applied fluorescence, absorbance, microscopy, MALDI-TOF spectrophotometry and other biochemical techniques to investigate the interaction between Tau and Baicalein *in vitro*. We found the aggregation inhibitory properties of Baicalein for the repeat Tau. Overall, the potential of Baicalein in dissolving the preformed Tau oligomers as well as mature fibrils can be of utmost importance in therapeutics for Alzheimer’s disease.

## Introduction

Alzheimer’s Disease is a neurodegenerative disorder characterized by memory loss and progressive dementia. The cytosolic Tau aggregates and the extracellular amyloid beta plaques are the typical traits of AD ^*1–3*^. The therapeutics for AD is being designed to act on both the targets in a discreet as well as integrated manner. The efficacy of the drugs designed seems to reduce as the level of trial increases ^*4, 5*^. The accumulation of cytosolic neurofibrillary tangles mainly composed of Tau PHFs ^*6*^ leads to neuronal degeneration. Along with microtubule stabilization Tau has varied roles such as neurite outgrowth, intracellular trafficking *etc.* that are hampered in AD due to elevated Tau levels ^*7, 8*^. All these neuronal abnormalities and degeneration due to Tau dysfunction and aggregation necessitates screening for molecules to inhibit Tau aggregation as well as to dissolve the preformed aggregates.

Tau aggregation in AD succeeds *via* several intermediates like oligomers, proto-fibrils, paired helical filaments formation which when untreated forms neurofibrillary tangles ^*9–11*^. Hence, these intermediates are being targeted for therapeutics in AD. The oligomer species are found to be more toxic than the filamentous aggregates ^*12*^ and hence Tau oligomers are gaining much attention as a therapeutic target in AD. Several molecules belonging to different classes like synthetic molecules, natural compounds, peptide inhibitors *etc.* have found to be potent against Tau aggregation ^*13–17*^. In recent time, methylene blue was found to be a potent molecule in AD ^*18–20*^ but failed to show effects on cognitive abilities in a transgenic mouse model of AD ^*21*^. On the other hand, the methylene blue derivatives MTC methylthioninium chloride (MTC); and its reduced form leucomethylthioninium salts (LMTX) has shown positive effects on Tau aggregation inhibition ^*22*^ by modifying the cysteine residues to the oxidized forms ^*23*^. Other approaches to inhibit Tau aggregation include targeting PTMs of Tau. Accordingly, kinase inhibitors or phosphatase activators have been screened for these molecules but failed to show potency due to low specificity and less efficiency. Since the ultimate effect of Tau aggregation is microtubule destabilization an alternative strategy for designing of therapeutics in AD is to screen or synthesize the microtubule stabilizing compounds ^*24–26*^. The natural compounds having varied targets in AD have also shown to have a potential as therapeutics in AD like curcumin, paclitaxel, geldanamycin *etc.*^*27*^. Azaphilones, a class of fungal metabolites have shown the aggregation inhibition as well as disaggregation activity for Tau aggregates ^*28*^. The secondary metabolites of *Aspergillus nidulans* like 2,ω-Dihydroxyemodin, Asperthecin, and Asperbenzaldehyde having aromatic rings are potent Tau aggregation inhibitors ^*29*^. Oleocanthal, derived from extra virgin olive oil have shown inhibition of Tau aggregation by covalently modifying the protein at its lysine residues ^*30*^. EGCG is an active component of green tea and has been shown to interfere with the aggregation of amyloid beta, α- synuclein as well as Tau in *in vitro* studies ^*31, 32*^. Baicalein is a polyphenol compound belonging to the class of flavonoids isolated from the Chinese herb *Scutellaria baicalensis*. Flavonoids are a rich source of anti-oxidants and are known to play a neuroprotective role ^*33*^. Baicalein has been known to exert anti-allergic, anti-carcinogenic anti-inflammatory as well as cardio protective properties. It has also shown a potential for inhibiting aggregation of proteins involved in neurodegenerative diseases. Baicalein not only prevents fibrillization of α-synuclein but also disaggregates the preformed fibrils involved in Parkinson’s disease ^*34*^. In AD transgenic mouse model, Baicalein has shown to reduce the amyloid-β accumulation and enhance the processing of amyloid precursor protein *via* α-secretases thus leading to non-amyloidogenic processing of APP ^*35*^. In this study, we report that Baicalein shows a dual property of inhibiting aggregation as well as dissolving the pre-formed fibrils of repeat domain of Tau in *in vitro* conditions at sub micro-molar concentrations. To explore the putative binding site and mode of Baicalein binding with Tau, the molecular docking and simulations were done with the hexapeptide repeat region. Only this region is amenable to predict the reliable model from the presently available structure information. The dynamics simulation on Tau filament reported previously showed the aggregation mechanism due to hyperphosphorylation ^*36*^. This study would reveal nature of interaction and mechanism of Baicalein in inhibition of Tau aggregates formation.

## Experimental Procedures

### Materials or Chemicals

Baicalein, ThS, ANS, BES, sinapinic acid BCA reagent and MTT were purchased from Sigma. Heparin, NaCl and sodium azide were purchased from MP Biomedicals. DTT and protease inhibitor cocktail were purchased from Calbiochem and Roche respectively. ECL reagent (32132) and antibodies anti-oligomer A11 rabbit polyclonal (AHB0052) and goat anti-rabbit HRP conjugated (A16110) were purchased from Thermo Fisher Scientific. The cell culture reagents DMEM F12, Fetal Bovine Serum (FBS), Penicillin-Streptomycin (Penstrep), L-glutamine, trypsin-EDTA, 1X phosphate buffered saline (PBS) were purchased from Invitrogen. 10 mM and 1 mM stock of Baicalein was prepared in ethanol. 12.5 mM mother stock of ThS was prepared in 1:1 ethanol: miliQ water, which was further, diluted to 200 μM stock in filtered miliQ water. ANS was prepared as 10 mM stock in filtered miliQ.

### Tau purification

The protein purification was carried out as previously described ^*37*^. In brief, the cell pellets obtained after induction of protein expression were homogenized under high pressure (15,000 psi) in a microfluidics device for 15 minutes. The obtained lysate was heated at 90°C for 15 minutes after addition of 0.5 M NaCl and 5 mM DTT. The heated lysate was then cooled and centrifuged at 40,000 rpm for 50 minutes. The supernatant was collected and dialyzed in Sepharose A buffer overnight. The obtained dialyzed sample was then subjected to a second round of ultracentrifugation and the supernatant was loaded onto the cation exchange column (Sepharose fast flow GE healthcare) for further purification. The bound protein was eluted using an ionic gradient. The eluted proteins were pooled and concentrated for size exclusion chromatography (16/600 Superdex 75pg GE healthcare). The Tau concentration was measured using BCA method.

### Molecular modeling

To identify templates for homology model building, a similarity search using Basic Local Alignment Search Tool (BLAST) ^*38*^ algorithm was performed against the Protein Data Bank (PDB), to identify high-resolution crystal structures of homologous proteins 63, 64. The sequence identity cut off was set to ≥ 30% (E-value cut off =1). Homology modeling of Tau K18 was then carried out using Modeller 9.16 ^*39*^ by taking structures homologous to the target proteins as templates, in order to study their structural features, binding mode and affinity with the substrates.

### Model validation and refinement

The initial models obtained, were evaluated for the stereochemical quality of the protein backbone and side chains using PROCHECK and RAMPAGE ^*40, 41*^. ERRAT server checked the environments of the atoms in the protein model ^*42*^. Errors in the model structures were also checked with ProSA server ^*43*^. After model validation, initial models were refined using impref minimization of protein preparation wizard and Impact 5.8 minimization ^*44*^. These energy minimized final models were further used for the binding studies with Baicalein.

### Ligand-protein preparation and docking studies

The ligand molecules considered in the present study were downloaded from ZINC compound database in the mol2 format. The protein and ligands were prepared first before proceeding with the docking studies. The water molecules and other hetero atom groups were removed from the protein structures using protein preparation utility of Maestro. Hydrogens were added subsequently to carry out restrained minimization of the models. The minimization was done using impref utility of Maestro in which the heavy atoms were restrained such that the strains generated upon protonation could be relieved. The root mean square deviation (RMSD) of the atomic displacement for terminating the minimization was set as 0.3 A°. Similarly, ligands were refined with the help of LigPrep 2.5 to define their charged state and enumerate their stereoisomers. The processed receptors and ligands were further used for the docking studies using Glide 5.8 ^*45*^. Results from fluorescence microscopy and MALDI-TOF spectrophotometry, indicated that certain flavonoids interact with the Tau protein, influencing their aggregation and disaggregation propensities. Thereby one such flavonoid originally isolated from the roots of *Scutellaria baicalensis* and *Scutellaria lateriflora*, believed to enhance liver health, Baicalein was docked in the binding site cavity of the prepared receptor molecule, aka the protein. A Sitemap analysis ^*46*^ was performed on the prepared receptor molecule, to identify the probable binding sites for our ligands of interest, since no prior information was available regarding the same. Next, selecting any of the Sitemap points obtained above generated grids. Flexible ligand docking was carried out using the standard precision option.

A total of 6 poses with the respective ligand and different sites were generated and scored on the basis of their docking score, glide score and E-model values. The hydrogen bond interactions between the protein and ligands were visualized using PyMOL.

### Molecular dynamics simulations

The docked complexes were prepared first using protein preparation wizard and then subjected to molecular dynamics simulations for a time scale of 15 nanosecond (ns) using Desmond 3.1 of Maestro ^*47*^. OPLS2005 force field was applied on docked complexes placed in the center of the orthorhombic box solvated in water. Protein was immersed in orthorhombic water box of SPC water model. Total negative charges on the docked structures were balanced by suitable number of counter-ions to make the whole system neutral (10 Cl-ions). The system was initially energy minimized for maximum 2000 iterations of the steepest descent (500 steps) and the limited memory Broyden-Fletcher-Goldfarb-Shanno (BFGS) algorithm with a convergence threshold of 1.0kcal/mol/Å. The short and long-range Coulombic interactions were handled by Cutoff and Smooth particle mesh Ewald method with a cut off radius of 9.0Å and Ewald tolerance of 1e_09. Periodic boundary conditions were applied in all three directions. The relaxed system was simulated for 10ns with a time step of 2.0 femtosecond (fs), NPT ensemble using a Berendsen thermostat at 300K temperature and atmospheric pressure of 1bar. The energies and trajectories were recorded after every 2.0 picosecond (ps). The energies and RMSD of the complex in each trajectory were monitored with respect to simulation time. The C-alpha atom root mean square fluctuation (RMSF) of each residue was analyzed. The intermolecular interactions between the target and substrate were assessed to check the stability of the complexes.

### Tau aggregation inhibition assay

Tau aggregation inhibition assay was performed with a few modifications in the already described protocol. In brief, the soluble Tau protein was centrifuged at 60,000 rpm for 1 hour (Optima Max XP Beckman Coulter) to remove aggregates present if any and 20 μM of soluble Tau protein was incubated in 20 mM BES buffer pH 7.4 with 5 μM of Heparin 17,500 Da in presence of 25 mM NaCl, 1mM DTT, protease inhibitor cocktail, 0.01% Sodium azide, and different concentrations of Baicalein ranging from 0 to 500 μM. The reaction mixtures were incubated at 37°C.

### Soluble protein assay

The soluble protein of repeat Tau was centrifuged as mentioned above prior setting up the reaction. 20 μM of each protein was incubated with 25 and 100 μM of Baicalein in absence of heparin. The other reaction mixture composition was maintained as mentioned above. A positive and negative control was kept with and without addition of heparin respectively. The reaction mixtures were incubated at 37°C.

### UV-Visible Spectrophotometric analysis and determination of the dissociation constant (Kd) of Tau-Baicalein interaction

50 μM Baicalein in 20 mM BES buffer was placed in a cuvette and an absorption spectrum was recorded in 230-450 nm range using Jasco V-530 spectrophotometer. Aliquots of repeat Tau were added to the cuvette in 3.5, 7, 14, 28, 42, 56, 70, 84, 98 μM concentration and respective spectra were recorded for each addition. The total volume of repeat Tau added was less than 100 μl so that there is no dilution effect.

### Repeat Tau disaggregation assay for oligomers and fibrils

The effect of Baicalein on the dissolution of Tau aggregates was carried out by incubating the preformed PHFs with different concentrations of Baicalein at 37°C. For disaggregation of repeat Tau oligomers, Baicalein was added after incubating the reaction mixture for 4 hours and for fibril disaggregation Baicalein was added at 60 hours. The Tau aggregates were prepared in 20 mM BES buffer pH 7.4 with 20 μM conc. of Tau and 5 μM final conc. of heparin. ThS and ANS fluorescence assays monitored the disaggregation. 20 μl samples from different time points were loaded onto 17% SDS-PAGE and analyzed using Bio-Rad Image Lab software.

### ThS fluorescence assay

ThS fluorescence assay monitored Tau aggregates formation as described. The 5 μl of 20 μM reaction mixtures were diluted in 50 mM ammonium acetate pH 7.0 containing 8 μM of ThS to a final volume of 50 μl. The ratio of protein to ThS was maintained as 1:4 for the fluorescence measurement. Fluorescence measurements were carried out using Tecan Infinite 200 Pro series plate reader. The excitation and emission wavelengths were set at 440 nm and 521 nm respectively. The plate was given a quick shaking for 10 seconds before the measurement. The readings were carried out in triplicates for each sample at 25°C. Initial readings were taken within an interval of 1 hours till 6 hours and later readings were continued at an interval of 12 hours each. The fluorescence was normalized for background by subtracting the buffer blank fluorescence.

### ANS fluorescence assay

The level of protein hydrophobicity was measured by ANS fluorescence. The 5 μl of 20 μM reaction mixtures were diluted in 50 mM ammonium acetate pH 7.0 containing 40 μM of ANS to a final volume of 50 μl. The ratio of protein to ANS was maintained as 1:20 for the fluorescence measurement. The excitation was carried out at 390 nm and the emission data was collected at 475 nm. Each sample was read in triplicate and subtracting from respective blanks eliminated the background fluorescence. All measurements were carried out in Tecan Infinite 200 Pro series plate reader at 25°C. The time points were followed in a similar way as ThS fluorescence.

### SDS-PAGE analysis

Samples left after fluorescence assay were collected and analyzed by SDS-PAGE for visual detection of aggregation inhibition on 17% polyacrylamide gels with different time points. The SDS-PAGE gel quantification was carried out using Image Lab software (Bio-Rad). The quantification for complete individual lanes was carried out using the software and the obtained intensities were plotted in the form of the bar graphs.

### Filter Trap assay

The detection of repeat Tau oligomers induced by Baicalein and separated by SEC was carried out by filter trap assay using A11 anti-oligomers rabbit polyclonal antibody. The fractions obtained after SEC were blotted onto Protran 0.45 μm membrane (Amersham Biosciences) using Minifold Dot-Blot System (Amersham Biosciences). The membrane was blocked with 10% milk in PBST (0.01% Tween 20) for one hour at room temperature. The primary antibody A11 was diluted to 1:2000 and the blot was incubated with it overnight at 4°C. After washing the unbound antibody thrice with PBST, the blot was incubated with secondary goat anti-rabbit antibody conjugated to HRP for one hour at room temperature in 1:10000 dilutions. The unbound antibody was washed thrice with PBST. The blot was developed using ECL-Plus reagent and the chemiluminescence was detected on Amersham Imager 600 and quantified using Image Quant TL software.

### Circular Dichroism Spectroscopy

CD spectroscopy was performed for determination of conformational changes occurring during the aggregation inhibition process using Jasco J-815 CD spectrometer in a cuvette with a path length of 1 mm under nitrogen atmosphere. The scan was carried out with bandwidth 1 nm, scan speed 100 nm/min; scan range 190 to 250 nm, and an average of 5 acquisitions were taken. In each experiment, measurements were done at 25°C. The buffer baseline was set with sodium phosphate buffer, pH 6.8. The concentration was maintained 1 μM for repeat Tau for all CD measurements.

### Electron microscopy measurements

Transmission electron microscopy was carried out to qualitatively determine the Tau filaments formation in presence and absence of Baicalein. 2 μM of the reaction mixtures were applied to 400 carbon coated copper grids for 45 seconds followed by 2 washes with filtered MilliQ water for 45 seconds each. The samples were negatively stained for 1 minute using 2% uranyl acetate. The dried grids were analyzed using Tecnai G2 20 S-Twin transmission electron microscope.

### Size Exclusion chromatography for Tau Oligomers

Size Exclusion chromatography was carried out for repeat Tau in presence of Baicalein at different time points in order to characterize the effect of Baicalein on the tendency of oligomerization. Repeat Tau samples were prepared by incubating the protein in 20 μM concentration in 20 mM BES buffer, pH 7.4 with or without Baicalein in 200 μM final concentration. Heparin was added in 1:4 ratios as an inducer of aggregation. Baicalein was added to the reaction mixture after 4 hours of incubation at 37°C. The initial incubation was provided for the preparation of lower order oligomers. The time point at which Baicalein is added is considered as 0 hour. The protein sample was filtered prior to loading on to Superdex 75 10/300 GL column (GE). BSA and Lysozyme were loaded as standards in 3 mg/ml concentration and soluble repeat Tau was loaded as a control. The separation was carried out in PBS buffer. Samples at 0, 6, 12, 24 and 60 hours of incubation were subjected to size exclusion chromatography.

### MALDI-TOF analysis of Baicalein modified Tau

Repeat Tau protein (1 mg/ml) was incubated with different concentrations of Baicalein (5, 10, 25, 50, 100, 200, 500 μM) for 1 hour and 12 hours at 37°C respectively. The samples were then diluted 1:20 in Sinapic acid and spotted on the MALDI plate and analyzed using ABSCIEX 4800 MALDI-TOF analyzer.

### Cell culture and toxicity assays

The neuroblastoma N2a cells were seeded in 96 well culture plates at the cell density of 25,000 cells/well. The cells were grown in DMEM F12 media containing 0.5% FBS and antibiotic (Penstrep) for 24 hours. For studying the toxicity of Baicalein induced oligomers 10 μL from the SEC fractions was added to cells/well and incubated for 24 hours. 10 μl of 5 mg/ml MTT (Methylthiazolyldiphenyl-tetrazolium bromide) was added to each well and incubated for four hours at 37 °C. The formazan crystals formed after reduction of MTT by viable cells enzymes were dissolved in 100 μl of 100% DMSO. The purple colour developed was read at 570 nm in TECAN plate reader.

### Statistical analysis

Sigma Plot 10.2 carried all the statistical analyses out using unpaired T-test. The error bars represent mean ±SD values. 95% confidence intervals were maintained for the analyses.

## Results

### Baicalein binds to Tau *in vitro*

Tau protein is one of the major microtubule-associated proteins in neuronal axons that mainly functions to stabilize and assemble microtubules. Tau is highly soluble protein and adopts a natively unfolded structure in solution ^*48*^. The repeat domain of Tau plus the proline rich flanking regions confer the property of microtubule binding and assembly (Fig. 1A), where the repeat domain of Tau represents the core of the PHFs. The chemical structure of Baicalein is depicted in (Fig. 1B). Baicalein shows characteristic λ_max_ at two wavelengths 270 nm and 362 nm (Fig S1B, C). Upon titrating with repeat Tau from 0-98 μM, the peak at 362 nm was found to decrease in intensity indicating the binding of Baicalein (Fig. 1D, S1C). The maxima at 362 nm plotted against increasing Tau concentration (0-98 μM) shows a gradual decrease in absorbance intensity (Fig. 1E). The dissociation constant (K_d_) for the binding of Tau with Baicalein was found to be 3.5 μM by using the following formula:

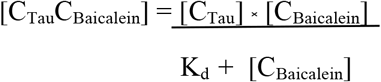

Where [C_Tau_] is the initial Tau concentration, [C_Baicalein_] is the free Baicalein concentration, [C_Tau_C_Baicalein_] is the concentration of Tau-Baicalein complex and K_d_ is the dissociation constant ^*49–51*^

**Figure 1.**
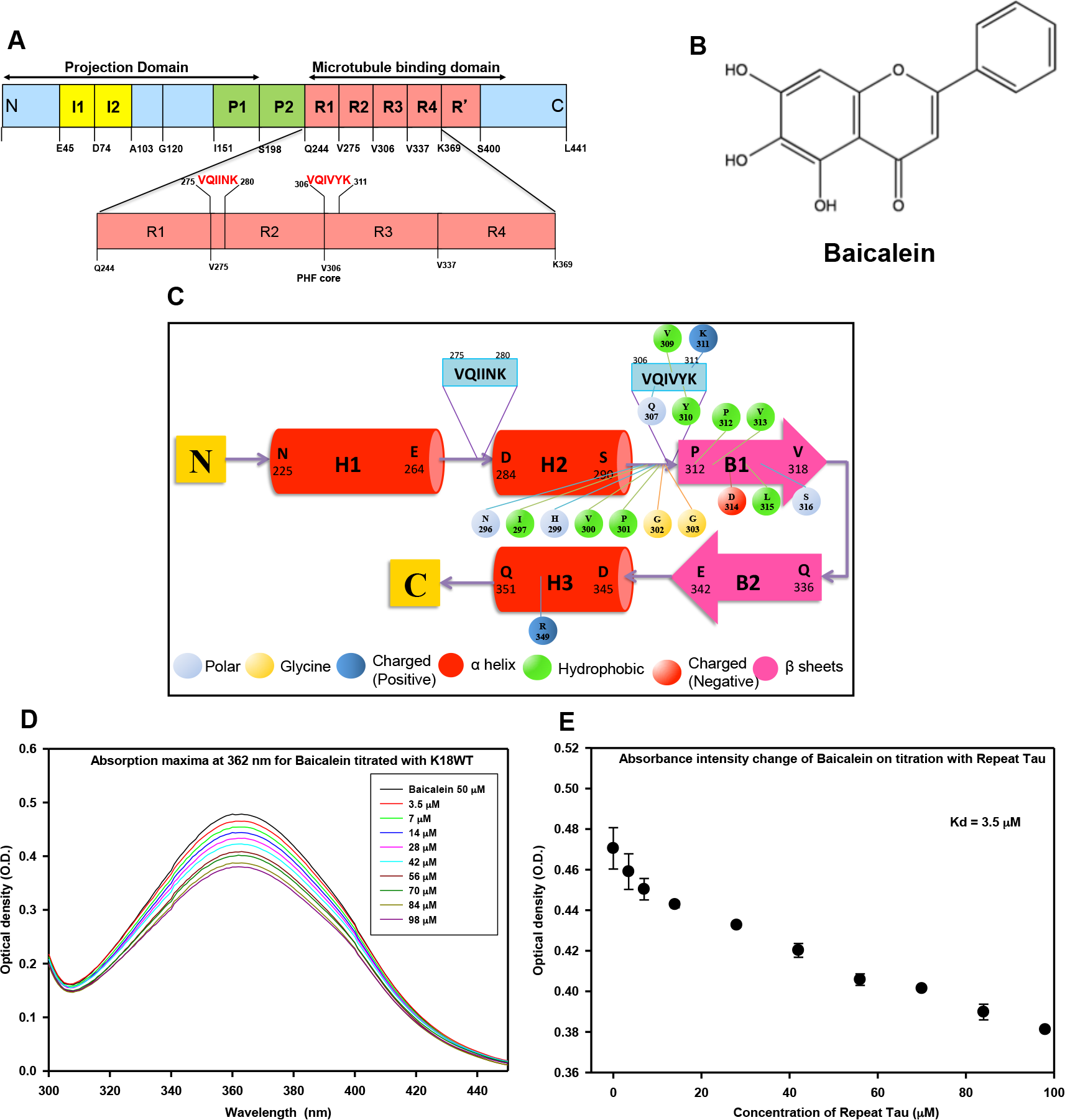
The interaction of Tau with Baicalein. A) Bar diagram showing full-length and repeat Tau. The figure shows longest isoform of human Tau with 441 amino acids. The yellow blocks represent two inserts of 29 amino acids each at the N-terminal region followed by the polyproline rich region, which gives a transient helix structure. The 4 red boxes represent the four repeats of 30 amino acids each. The K18 construct comprises of only the repeat domain of Tau, which forms the core of Paired Helical Filaments. B) Chemical structure of the flavonoid Baicalein. C) A schematic diagram of the repeat Tau model secondary structure depicting three predicted α helices (H1, H2, and H3) and two predicted β sheets (B1, B2) with two distinct and characteristic hexa-peptide regions ^275^VQIINK^280^ and ^306^VQIVYK^311^. The predicted ligand interacting residues, color coded according to their type of interaction is also shown. D) Figure shows decrease in the absorbance intensity at 362 nm upon increasing the protein concentration against Baicalein. E) The absorbance intensity at 362 nm obtained for different titrations when plotted against respective concentrations gives a plot representing the binding affinity of Tau with Baicalein.

### Molecular Models of Tau protein

The molecular structures of repeat Tau were modeled, in order to ascertain the putative binding site of Baicalein and the role of the conserved hexapeptide repeat regions ^275^VQIINK^281^ and ^306^VQIVYK^311^ in the interactions with Baicalein (Fig. 1C). It is very challenging to predict the full-length Tau model due to its disordered nature of the regions in both N- and C-terminal. But the hexapeptide repeat region forms the stable secondary structure as reported by many biochemical studies and structural information ^*52–54*^. A BLAST search against PDB identified homologous structures, to be used as templates to build the homology models of this region. Structures of Tau (267-312) bound to microtubules (2MZ7) and shared an identity of 100% and query coverage of 92% with the target sequence, and were thus used for modeling studies.

The initial models were evaluated for their stereochemical parameters. More than 98% of all residues were in the allowed regions of Ramachandran plot. The model refinement phase involved preprocessing the initial models by adding hydrogen’s, assigning bond order, and filling missing loops and side chains. Next, the models were subjected to restrained minimization by applying the constraint to converge the non-hydrogen atoms to an RMSD of 0.3 using OPLS 2005 force field. Subsequently, the models were subjected to 500 steps of steepest descent energy minimization followed by 1000 steps of conjugate gradient energy minimization using the same force field. These energy-minimized models were used for docking and molecular dynamics studies.

### Molecular docking of repeat Tau and Baicalein

The docking studies of Baicalein with repeat Tau model generated the best pose with glide score and E-model values of − 5.06 and − 44.365 respectively. The O_1_ atom of Baicalein interacts with the NH_2_ group of Asn 265 and the NH group of the imidazole ring of His 268. The OH groups involving O_3_ and O_4_ atoms of Baicalein, interacts with the carboxyl O^−^ atom of Glu 338, forming hydrogen bonds in the process (Fig. 2A). While the two -OH groups (O_3_ and O_4_) have an interaction distance of 2.70 Å, 2.0 Å and 1.80 Å respectively, the interactions with the amine group stand at a distance of 2.30 Å for Asn 265. Further hydrophobic interaction was observed with residues Tyr 310 and Val 313, of which one (Tyr 310), interestingly, belongs to the hexapeptide repeat ^306^VQIVYK^311^ and the other (Val 313) is a flanking residue to the same repeat, a prime area of interest in our study. Another crucial residue His 299 is predicted to have an important interaction with Baicalein via water bridges, which may be involved in the probable mechanism of Tau disaggregation in presence of Baicalein.

**Figure 2.**
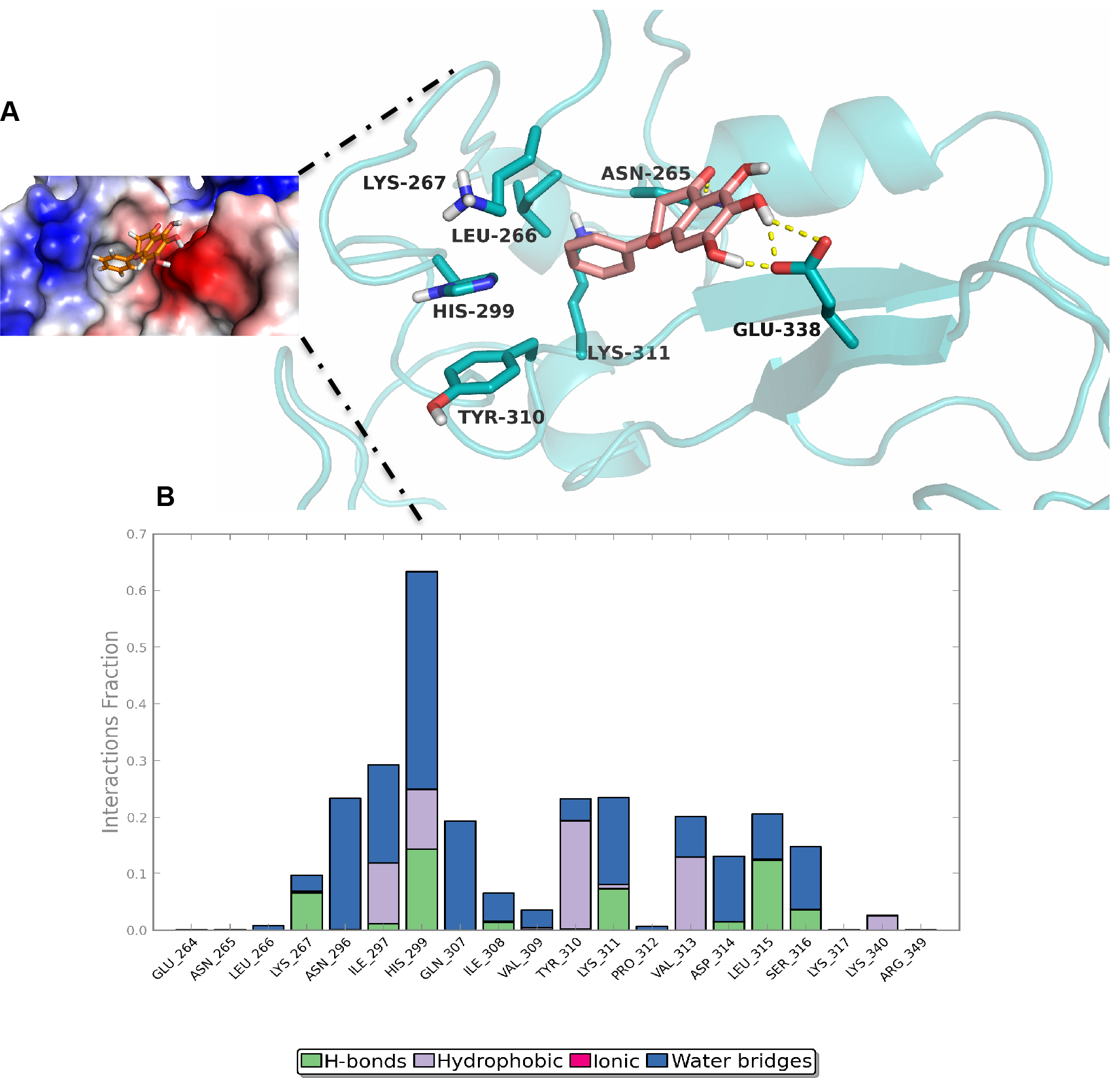
Docking and Simulation of Tau with Baicalein. A) A detailed snapshot of the ligand-binding pocket of Baicalein in our proposed model of repeat Tau including the interacting residues as indicated and the corresponding surface represented in terms of electrostatic potential. B) A bar diagram depicting the extent of interaction over the course of 15 ns simulation using the same docked structure which indicates that the residues mentioned are majorly involved in the binding and probable biochemistry of the ligand binding to Tau. The protein-ligand interactions or contacts are categorized into four types: Hydrogen Bonds, Hydrophobic, Ionic and Water Bridges.

### Molecular dynamics simulation of repeat Tau and Baicalein

The Baicalein docked complex was subjected to a simulation of 15ns time duration to analyze the stability of the interaction and other structural changes. The RMSD graph was plotted by assigning time in ns on the X-axis and RMSD values of the Cα atoms of the protein and ligand in /Å on the left and right Y-axes respectively (Fig. S2). The graph and the snapshots at different time intervals of simulation indicated that the RMSD of Cα atoms of the generated structures over the complete trajectory is quite stable with minor repositioning of Baicalein with respect to the hexapeptide repeat region ^306^VQIVYK^311^. The Cα RMSD between the initial and final conformation after the simulation of docked complex was ~10 Å indicated the considerable conformational change upon binding. During the course of the simulation, a few residues namely Asn 296, Ile 297, His 299, Gln 307, Tyr 310, Lys 311, Val 313 and Leu 315 out of which three residues belong to the hexapeptide region mentioned above, were found to form an increasingly high number of interactions with Baicalein, indicating that these may be crucial residues to stabilize the complex (Fig. 2B, S3).

### Baicalein inhibits Tau aggregation *in vitro*

Tau is a natively unfolded protein with an overall positive charge. This property makes it possible to aggregate Tau *in vitro* by using polyanionic co-factors like heparin. Heparin is a polymer of glycosaminoglycan monomers, which has a negative charge and induces aggregation of Tau in vitro ^*55, 56*^. This approach has been widely used to screen for the Tau aggregation inhibitors ^*18, 30, 57*^. The aggregation inhibition of the repeat Tau by Baicalein was determined using 0, 5, 25, 50, 100, 200, and 500 μM concentrations. The rate of Tau fibril formation was monitored by ThS fluorescence. The ThS fluorescence intensity decreased with increasing concentrations of Baicalein in the treated samples. The ThS fluorescence showed an initial increase in control as well as treated samples till 4 hours of incubation (Fig. 3A). As the time progressed, the treated samples showed decrease in fluorescence intensity but the control showed stagnant saturated fluorescence suggesting a concentration dependent inhibition of aggregation. The percent inhibition at the end of 60 hours of was 91% for higher concentrations of Baicalein (Fig. 3A, B). The inhibition was observed to occur at an early time point and the IC_50_ value for repeat Tau aggregation inhibition was found to be 35.8 μM of Baicalein (Fig. 3C). The pattern of rise and fall in fluorescence intensity for the treated samples suggest that Baicalein might be inducing initial oligomerization of Tau protein and inhibiting the further fibrillization by sequestering these oligomers.

**Figure 3.**
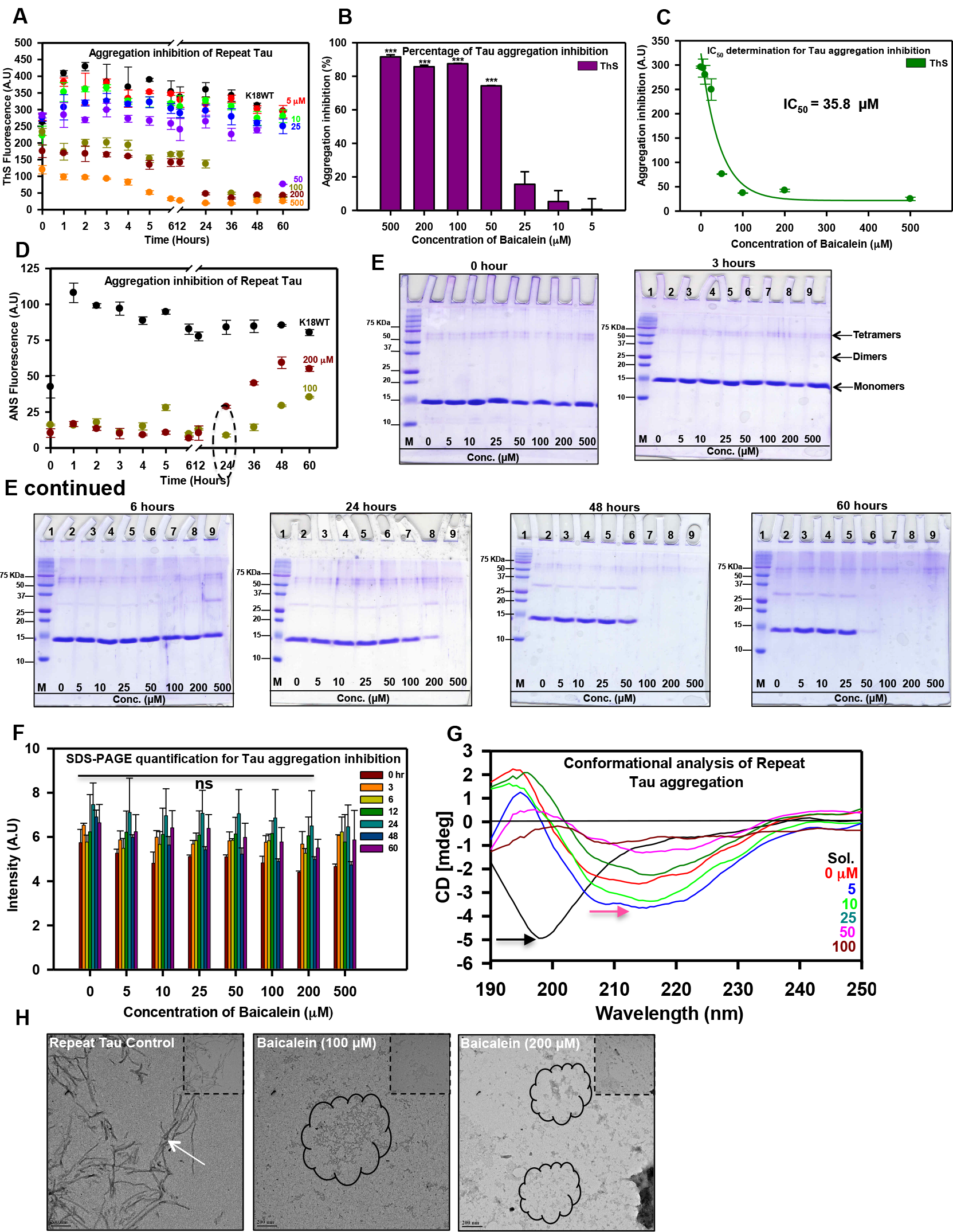
Tau aggregation inhibition by Baicalein. A) The ThS fluorescence show a concentration dependent decrease in intensity suggesting inhibition of repeat Tau aggregation by increasing dose of Baicalein. B) The maximum inhibition is shown by highest concentration (500 μM) of Baicalein showing around 90% aggregation inhibition. C) The IC_50_ value for repeat Tau aggregation inhibition was found to be 35.8 μM of Baicalein. D) The ANS fluorescence for the higher concentrations of Baicalein showing initial decrease and further rapid increase in fluorescence at 24 hours (dotted circle). E) The SDS-PAGE analysis at zero time point control as well as treated samples does not show presence of any higher order aggregates. With an increase in time there is an increase in higher order oligomers observed especially after 6 hours of incubation. At 24 hours, the 500 μM of Baicalein (lane 9) shows the presence of dimer as well as tetramer. With increase in incubation time (48 and 60 hours), the oligomers are seen in all treated samples with differential intensity. The further incubation reveals the disappearance of these oligomers in concentration dependent manner. F) Quantification for the SDS-PAGE at different time points and different Baicalein concentrations shows decrease in the aggregates at higher concentrations of Baicalein. G) The CD analysis of the repeat Tau aggregation shows a shift in the spectra (pink arrow) in control as well as treated samples as compared to soluble protein (black arrow) indicating a shift towards β-sheet structure. The Sol. represents for soluble protein. H) The electron micrographs show the presence of filamentous aggregates (white arrow) in control as opposed to amorphous aggregates (encircled black) in the Baicalein treated samples.

### Baicalein enhances Tau oligomerization in presence of heparin

The *in vitro* formation of PHFs in presence of heparin undergoes various intermittent stages ^*58*^, which results in changes in hydrophobicity of the Tau protein ^*59*^. One of the intermittent stages in PHFs formation is the formation of oligomeric species, which acts as a nucleation step to further enhance fibrillization. The primary studies with ThS indicated that Baicalein prevents Tau fibril formation. We further investigated the role of Baicalein in Tau oligomerization. As an intermediate in Tau fibrillization, Tau oligomers show enhanced hydrophobicity due to their partially folded characteristics. ANS is a dye, which emits enhanced fluorescence on interaction with partially, folded or exposed hydrophobic micro domains of the proteins.

Thus, to study the effect of Baicalein on Tau oligomerization and accompanied hydrophobicity changes, we monitored ANS fluorescence for the control as well as treated samples. ANS showed an initial decrease in fluorescence and then the rapid increase before becoming stagnant (Fig. S4). This effect was evident at the earlier time points in the higher concentrations of Baicalein (Fig 3D). This behavior of ANS suggests that Baicalein might be involved in initial Tau oligomer formation and sequestration.

### SDS-PAGE and Immunoblot analysis reveals the enhanced Tau oligomerization and sequestration by Baicalein

In order to confirm our hypothesis, we carried out SDS-PAGE analysis of the control as well as Baicalein treated samples at different time intervals. We examined 10 μl of all the reaction mixtures at zero time point to rule out the presence of any preformed higher order aggregates (Fig. 3E). At 6 hours of incubation, we observed the dimers and tetramers in the highest concentration of Baicalein (Fig. 3E, F) and these oligomers appear in all the concentrations of Baicalein treated samples in a time dependent manner. We observed a complete clearance of dimer and monomers in 500 μM Baicalein treatment after 12 hours of incubation but the band for tetramers still remains. At 24 hours, 200 μM of Baicalein treated Tau show complete conversion from soluble to higher order oligomers. It is to be noted that this is the exact time point when we see stable decrease in ThS fluorescence and an increase in ANS fluorescence (Fig. 3A, D). This seems to be the point where Baicalein sequesters the oligomers formed and inhibits further fibril formation.. These results indicate that Baicalein enhances the oligomerization of Tau and blocks them at an oligomeric stage without further leading to fibril formation.

### Baicalein binds and stabilizes partially folded Tau oligomers

Since Baicalein demonstrated the enhancement of Tau oligomerization and their sequestration, we proceeded further to elucidate the conformation of these oligomers by Circular dichroism spectroscopy. The soluble Tau protein is natively unfolded and showed CD spectra of typical random coil at 198 nm (Fig.3G). Tau on aggregation shows the structural transition from typical random coil to the β-sheet structure ^*54*^. Tau protein treated with Baicalein showed shift in the wavelength towards a partial β-structure (Fig. 3G) suggesting that Baicalein is able to bind and stabilize the partially folded Tau oligomers.

### Electron microscopy visualization of Baicalein induced Tau aggregation inhibition

The time dependent effect of Baicalein on Tau oligomerization and aggregation inhibition was observed by electron micrographs. The untreated sample showed the presence of long fibrillar structures, which are characteristics of Tau filamentous aggregates (Fig 3H). On the contrary, the 100 μM Baicalein treated Tau showed presence of only pieces of Tau filaments (Fig 3H). The electron micrograph observation supports our previous observations that Baicalein prevents Tau fibrillization.

### Baicalein alone does not cause Tau conformational change

As our results suggested that Baicalein could stabilize Tau oligomers having partial β-sheet structure, it was necessary to understand whether, Baicalein alone can cause the random coil to β-sheet transition of Tau protein. For this, we incubated repeat Tau with Baicalein in absence of heparin. We maintained a positive control (with heparin) and negative control (without heparin) for standard aggregation kinetics. The kinetics for both ThS and ANS did not show any increase in fluorescence for negative control as well as Baicalein treated samples (Fig. S6A, B). The positive control showed an increase in ThS and ANS fluorescence as expected. The SDS-PAGE analysis showed the presence of higher order structures in the positive control from 24 hours onwards (Fig. S6C). The sample treated with higher concentration of Baicalein (100 μM) showed presence of very faint higher order bands at a later time point for both the proteins (Fig. S6D). The fluorescence kinetics and the SDS-PAGE analysis clearly suggest that Baicalein might be able to induce Tau oligomerization but at a much slower pace. The structural analysis of these oligomers revealed that they do not show transition to β-sheet structure (Fig. S6E,) and do not form Tau fibrils (Fig. S6F). These results clearly suggest that Baicalein alone might be able to induce soluble Tau oligomers at a slower rate without causing any structural changes.

### Baicalein can disaggregate both oligomers and fibrils of repeat Tau

The effect of Baicalein was studied comparatively for disaggregation of repeat Tau oligomers and mature fibrils respectively. Baicalein was found to enhance and moderately induce oligomerization as well as sequester the oligomers. Analysis of disaggregation of repeat Tau using ThS fluorescence assay suggested that Baicalein is able to disaggregate preformed oligomers in a concentration dependent manner (Fig. 4A). This suggests the oligomers are sequestered into an aggregation incompetent form. The similar effect in ThS fluorescence was observed for repeat Tau fibrils disaggregation (Fig. 5A). The underlying mechanism of disaggregation may involve the breaking of aggregates into smaller fragments and then by inhibiting the reformation of higher order aggregates. There was up to 80% disaggregation activity observed for the sample incubated with highest concentration of Baicalein (500 μM) (Fig. 4B) in oligomer disaggregation while nearly 85% disaggregation was observed in case of fibril incubated with 500 μM Baicalein (Fig. 5B). It was observed that both ThS and ANS fluorescence was decreased upon addition of Baicalein in 4 hours (Fig. 4A, C) and 60 hours (Fig. 5A, C) incubated samples suggesting disaggregation and decreased hydrophobicity respectively. However, the ANS fluorescence increased after 12 hours in all the incubated samples due to charge interactions. The SDS-PAGE analysis showed that Baicalein could bind to the preformed oligomers and sequester them at this stage. The presence of a tetramer band in all the concentrations of Baicalein can be observed but the dimer as well as the monomer completely disappears (Fig. 4D). In case of fibril disaggregation study, Baicalein treated samples showed decrease in the population of oligomeric species as observed in SDS-PAGE for different time point samples (Fig. 5D). The fact that we observe the oligomers on the SDS-PAGE and increase in ANS fluorescence but a complete decrease in ThS fluorescence for the Baicalein treated oligomers as well as fibrils strongly suggests the role of Baicalein in sequestering the Tau oligomers. The SDS-PAGE quantification for repeat Tau oligomer (Fig. 4E) and fibril disaggregation (Fig. 5E) shows a gradual decrease in soluble repeat Tau in a concentration dependent manner. This is also supported by the electron micrographs (Fig. 4H and 5H), which shows fragility of Tau and absence of mature Tau filaments. Further, CD spectra analysis for repeat Tau samples treated with a varying range of Baicalein concentrations shows a shift towards the β-sheet structure in all concentrations (Fig. 4F, G, and Fig. 5F, G). This shows that Baicalein drives the soluble Tau towards formation of oligomers. It is also evident in size exclusion chromatography (SEC) for repeat Tau that treatment with Baicalein leads to formation of oligomers as seen in the SEC profiles for samples at 0, 6, 12, 24, 48 and 60 hours (Fig. 6A) as compared to soluble repeat Tau. Further the repeat Tau was incubated with Baicalein at 1:10 ratio showed presence oligomers while it was absent in the control aggregation sample at 60 hours, (Fig. 6B) suggesting enhancement of oligomerization in presence of Baicalein. There is an increase in the intensity of peak corresponding to repeat Tau oligomers with respect to time (Fig. 6C). This increase in the population of oligomers was also confirmed by probing the obtained fractions with A11 anti-oligomer antibody by filter trap assay (Fig. 6D). An increase in oligomeric population was observed with increase in incubation time, which is in conjunction with the prior results reiterating the role of Baicalein in inducing Tau oligomerization. To further elucidate the nature of these oligomers, we treated the neuro2a cells with the oligomer as well as soluble fractions obtained after SEC. The preformed oligomers treated with Baicalein at 0 hour were found to be more toxic as compared to 60 hours treated oligomers. The toxicity was reduced in presence of Baicalein though not 100% rescuing the viability of the cells (Fig. 6E).

**Figure 4.**
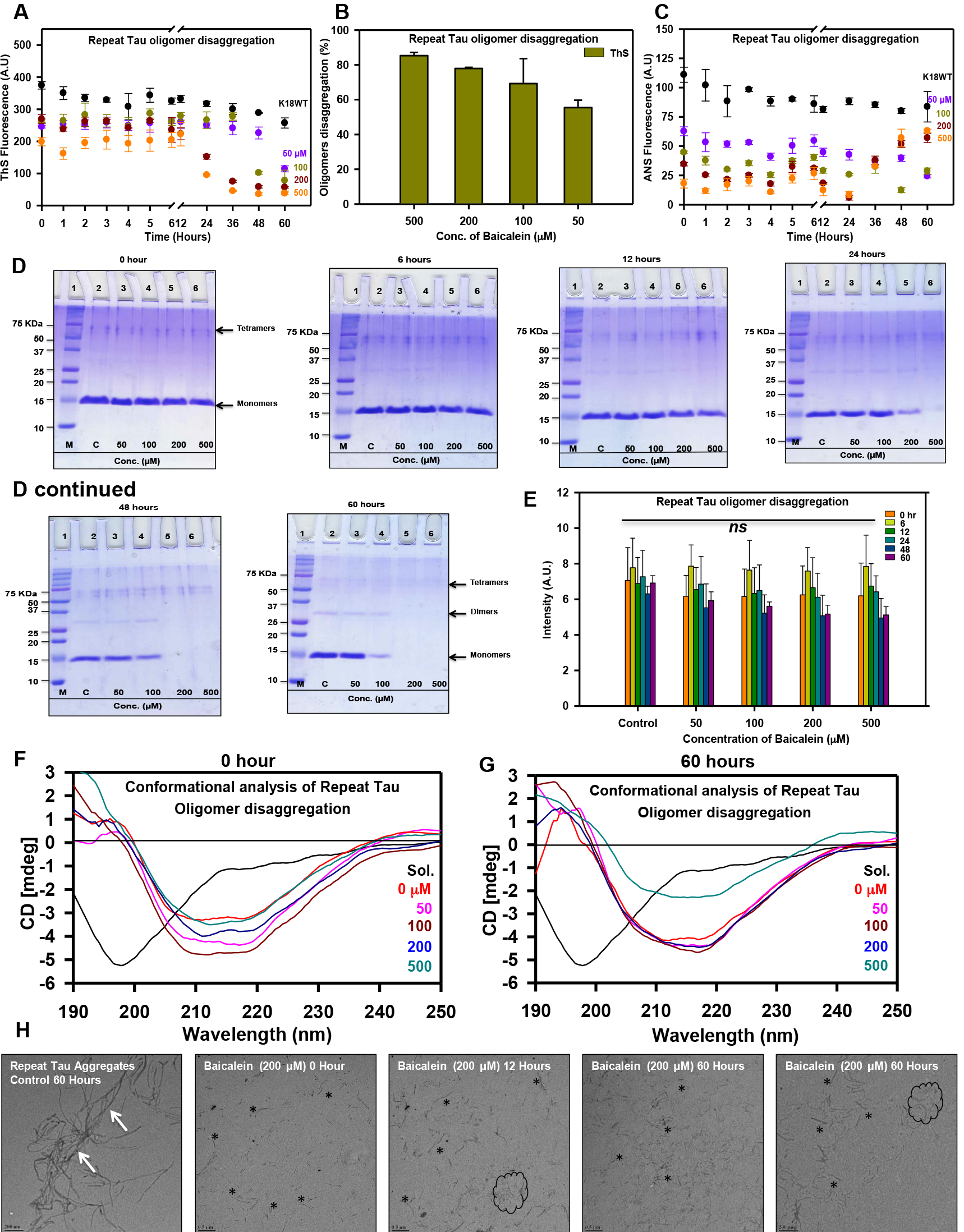
Disaggregation of repeat Tau oligomers by Baicalein. A) Formation of repeat Tau aggregates was monitored by ThS fluorescence after addition of Baicalein at different concentrations after 4 hours of incubation. ThS fluorescence shows a marked decrease in a concentration dependent manner. B) Percentage disaggregation of repeat Tau oligomers as compared to control at different Baicalein concentrations. The highest concentration of Baicalein (500 μM) shows 80% disaggregation while 50 μM Baicalein shows almost 60% disaggregation. C) ANS fluorescence assay for repeat Tau oligomers disaggregation on addition of different concentrations of Baicalein at 4 hours of incubation shows a prominent decrease in fluorescence after Baicalein addition in all the concentrations. There is an increase in fluorescence from 24 hours onwards indicating that at higher concentrations Baicalein can increase the hydrophobicity of protein aggregates. D) SDS-PAGE analysis shows a marked decrease in the monomeric form of repeat Tau, which are mostly entrapped as dimeric and tetrameric forms in presence of all different concentrations of Baicalein. E) Quantification of SDS-PAGE showing concentration dependent decrease in intensity for repeat Tau oligomers. F.G) The CD spectra for repeat Tau oligomer disaggregation samples at 0, and 60 hours of incubation with different Baicalein concentrations. In all the treated samples, the shift in the spectrum towards β-sheet structure can be observed (shown by pink arrow) as compared to soluble repeat Tau. The Sol. represents for soluble protein. H) Electron micrographs for repeat Tau at 0, 12 and 60 hours of incubation with 200 μM Baicalein show smaller broken filaments (black asterisks and encircled area) as compared to untreated control (white arrow).

**Figure 5.**
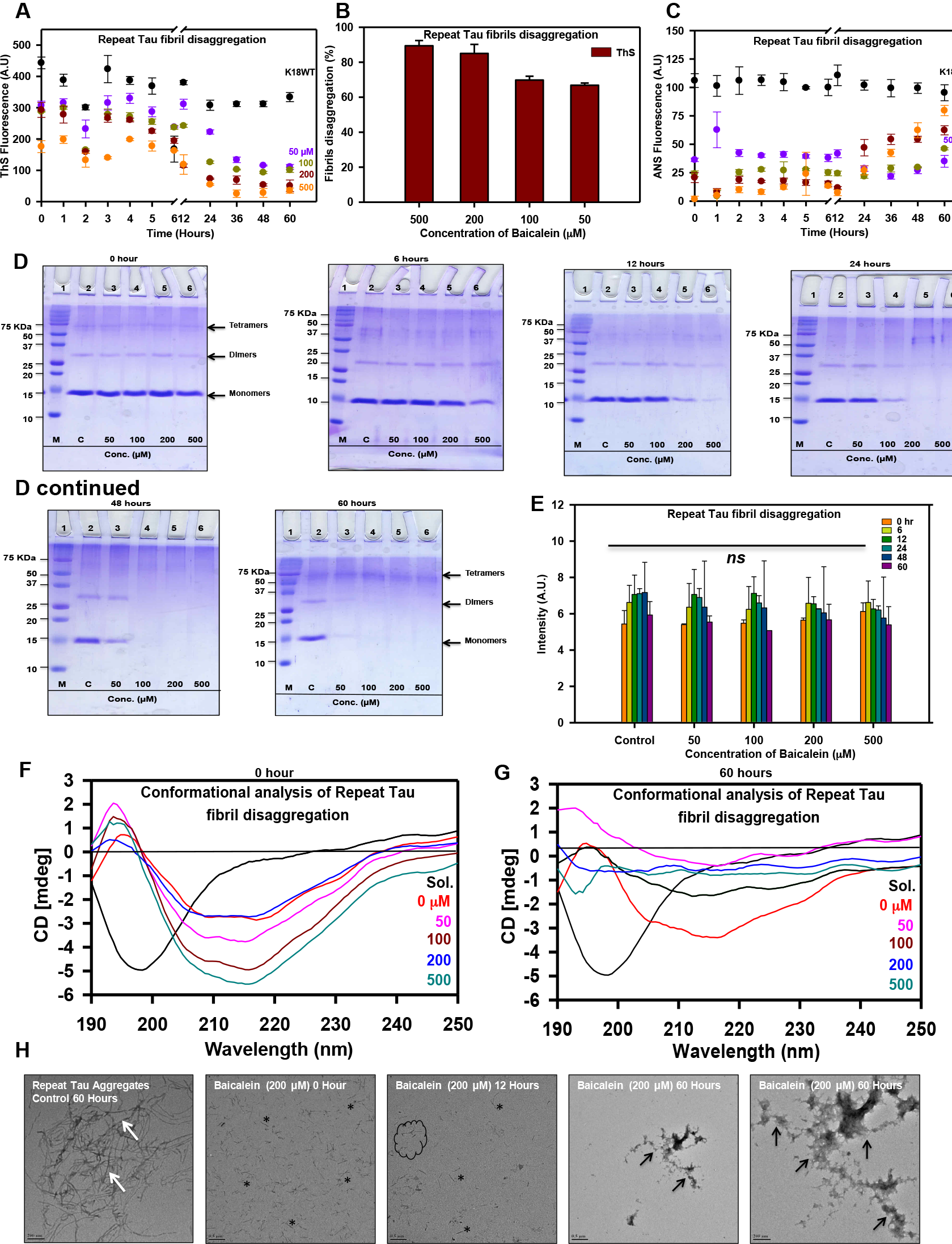
Disaggregation of repeat Tau fibrils by Baicalein. A) ThS fluorescence for repeat Tau fibril disaggregation shows a decrease with increase in Baicalein concentration and the time of incubation. B) The effect of Baicalein is concentration dependent as observed by percentage disaggregation, where, there is as much as 85% activity at the highest Baicalein concentration (500 μM) while nearly 60% activity at 50 μM concentration. C) The ANS fluorescence assay for repeat Tau fibrils disaggregation shows more prominent decrease in all concentrations of Baicalein. Furthermore, there is similar increase in ANS fluorescence after 12 hour of Baicalein addition. D) The SDS-PAGE analysis for different time point samples shows enhancement of repeat Tau dimer and tetramers until 12 hours after Baicalein addition. At 60 hours, the population of repeat Tau soluble protein as well as dimer is greatly reduced in all Baicalein concentrations while the tetrameric population is enhanced. E) Quantification of SDS-PAGE for samples with 50, 100, 200 and 500 μM of Baicalein was carried out for 0 hours to 60 hours of Baicalein addition. F, G) CD spectra for repeat Tau aggregates at different time points treated with Baicalein at different concentrations shows a shift in structure towards β sheet (shown by pink arrows) as compared to soluble repeat Tau. The Sol. represents for soluble protein. H) Electron micrographs for repeat Tau control and fibrils at different time points with 200 μM of Baicalein show disaggregation of fibrils into small pieces (black asterisks and encircled area) as compared to untreated control (white arrow).

**Figure 6.**
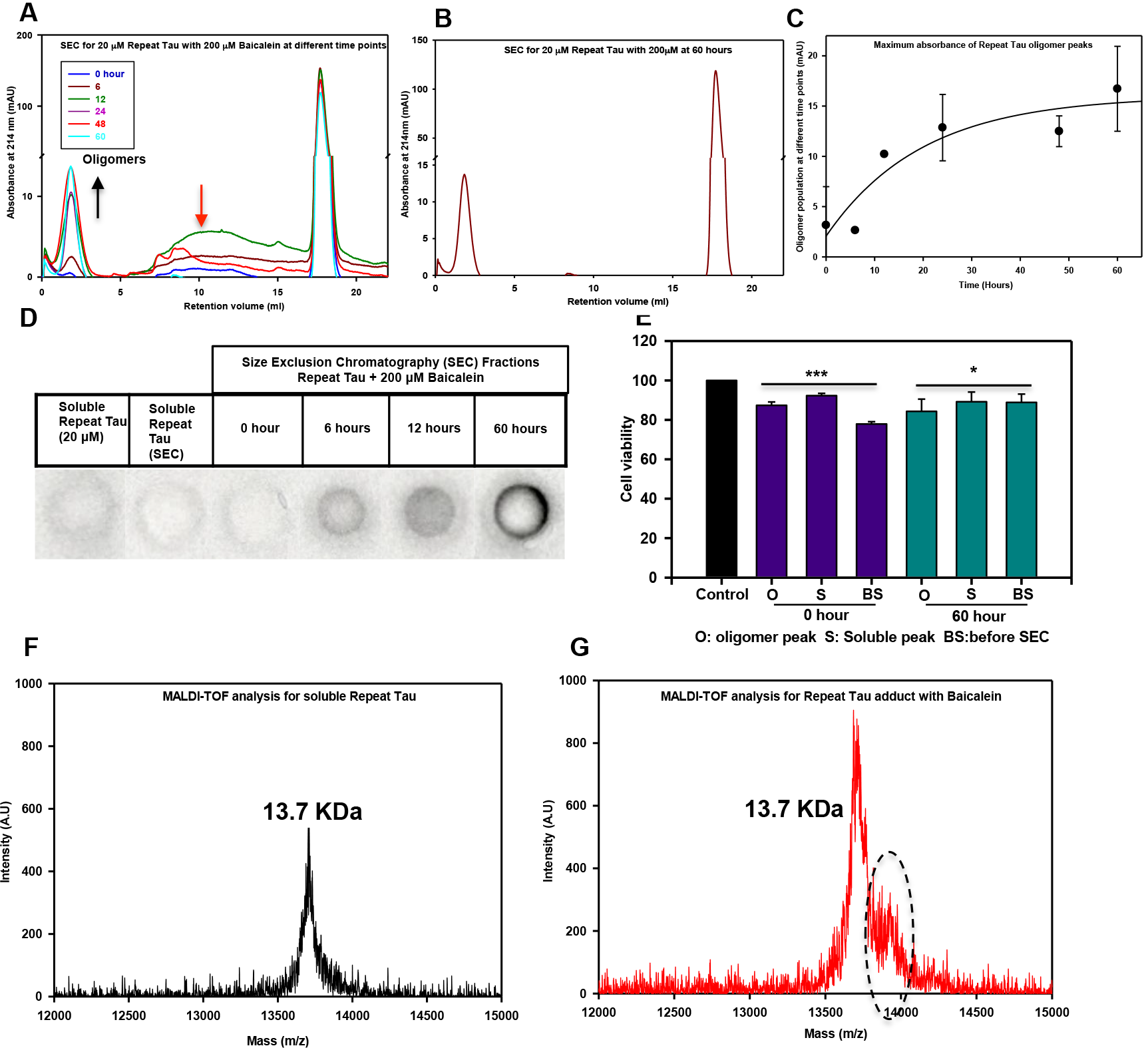
Size-exclusion chromatography for repeat Tau aggregates. A) Comparison of the chromatograms for repeat Tau incubated with Baicalein at different time points. Figure shows a decrease in dimeric population of repeat Tau (pink arrow) with subsequent increase in the Baicalein stabilized oligomers (black arrow). B) Chromatogram for repeat Tau incubated with Baicalein for 60 hours shows increase in oligomer population. C) Maximum absorbance obtained for the peaks for oligomers at different time points shows a gradual increase in Baicalein stabilized oligomers with time. D) The Baicalein induced repeat Tau oligomerization observed by SEC was confirmed by filter trap assay using A11 (anti-oligomers antibody). No significant signal was observed for the soluble repeat Tau before and after SEC confirming absence of oligomers in untreated sample. E) The oligomer as well as the soluble fractions obtained after the SEC of Baicalein treated repeat Tau did not show toxicity on N2a cells. F) The MALDI-TOF analysis of soluble repeat Tau showing 13.7 KDa molecular weight. G) The Baicalein modified Tau showing an adjacent peak of 13.9 KDa in addition to 13.7 KDa peak suggesting covalent modification of repeat Tau by Baicalein.

### Baicalein covalently modifies Tau protein as observed by MALDI-TOF

Baicalein showed oligomer inducing property for Tau in presence of heparin and sequestering these oligomers from further fibrillization. To understand the mechanism by which this occurs, MALDI-TOF analysis was carried out to study whether Baicalein can lead to covalent modification of Tau. The repeat Tau having a theoretical molecular weight of 13.7 KDa was incubated with a range of concentrations of Baicalein for 1 hour and 12 hours respectively. The samples incubated for 1 hour showed an additional peak of 13.9 KDa at higher concentration of Baicalein (Fig. 6F, S5D), which was not observed in untreated samples (Fig.6G, S5A) as well as in lower concentrations of Baicalein (Fig. S5B, C). But after 12 hours of incubation, all Baicalein treated samples showed a peak adjacent to the peak given by soluble Tau (13.7 KDa) of 13.9 KDa suggesting the covalent modification of repeat Tau by Baicalein (270 Da) (Fig. S5F, G, H) as opposed to untreated (Fig. S5E). Thus, Baicalein could be involved in covalently modifying Tau protein, thus, preventing it from aggregation.

## Discussion

Aggregation of Tau into insoluble aggregates inside the neuronal cells contributes to the pathogenesis of Alzheimer`s disease. The screening of compounds against Tau neurofibrillary tangles has yielded variety of potent molecules ^*60–62*^. Tau pathology undergoes a formation of series of intermittent species and the feasibility of their formation depends on both physiological concentration of free Tau and ability to form aggregation competent conformations ^*63*^. The soluble oligomers of Tau are more toxic than the Tau fibrils ^*64*^ and hence the Tau oligomers are gaining more attention as therapeutic targets in AD ^*65*^.

The screening for Tau aggregation inhibitors has generally yielded compounds with ring structures, which might have differential interactions with the Tau monomer, Tau aggregates and their intermediates ^*14, 57*^. Our current study elucidates that the herbal compound Baicalein binds to specific hydrophobic patches of the repeat domain of Tau and inhibits the Tau aggregation by targeting Tau into soluble oligomers. Similar results have been observed for the structurally similar polyphenol altenusin wherein *in vitro* Tau pathology is inhibited by the polyphenol but the *in vivo* pathology in transgenic mice could not be rescued ^*66*^. The progress of Tau aggregation leads to formation of more β-sheet rich structure ^*54*^, which provides the stability to these aggregates. Tau aggregation inhibitors either act to weaken the cross beta structures by inserting their rings between the beta sheet stacks or modify the protein or its intermediate aggregates, thus decreasing its aggregation potency ^*67*^. We propose Baicalein belongs to the latter class of inhibitors as it binds and stabilizes the high molecular weight Tau oligomers, which are not capable of fibrillization. The ANS fluorescence clearly suggest the increased hydrophobicity of these sequestered molecules ^*59, 68*^. The MALDI analysis reveals the modification of Tau by Baicalein as it forms adduct with repeat Tau as evidenced by an increase in molecular weight of Tau with Baicalein.

In order to shed light on the binding site and the molecular nature of Baicalein interaction with the Tau monomer, we have performed the docking and simulation studies. As the full-length Tau model is impossible to predict due to the disordered nature of the protein, we could get reliable model for the structured regions of Tau only, which is hexapeptide repeat region (244-373). Our results demonstrated that the Baicalein binds to this region. The MD simulation demonstrated that the structure was stable and the nature of interaction is primarily mediated by hydrophobic interaction and later stabilized by the water-mediated hydrogen bonds. Also, the trajectory of complex simulation suggests the observable conformational change of this aggregation prone region of Tau after Baicalein binding. This conformational dynamicity may be due to the relatively flexible bond between C7-C4 of the Baicalein. This observation is in agreement with the structure of Baicalein as it has two adjacent -OH groups that might have the ability to modify the Tau oligomers and form high molecular weight aggregates, which are visible on SDS-PAGE. Similar mechanism has been postulated for Tau aggregation inhibition with respect to other flavonoids having adjacent -OH groups ^*69*^. Hence, it can be conceived that Baicalein belongs to the class of covalent inhibitor of aggregation. The other potent inhibitors of this class include Oleocanthal, an aldehyde by nature, which reacts with the epsilon amino group of lysine residues to form imines ^*70*^. The mechanism of other covalent inhibitors like cinnamaldehyde and asperbenzaldehyde ^*29*^, which are α, β unsaturated aldehydes, involves nucleophilic attack by the cysteine residues of Tau, In contrast, Baicalein a polyalcohol flavonoid can undergo oxidation to Quinone form, which is involved in electrophilic attack ^*71*^. The previous studies with aggregation inhibition of α-synuclein have also revealed the similar mechanism of inhibition wherein the oligomers of α-synuclein are prevented to undergo fibril formation in presence of Baicalein ^*34*^. Thus, any inhibitor, which can serve the function of targeting these nucleating species, can be of great therapeutic importance. The effect of Baicalein must be prior to the nucleation as its inhibitory effect on Tau aggregation can be observed early in the course of *in vitro* aggregation. The size exclusion chromatography results also support this hypothesis, as the Baicalein treated samples showed the disappearance of a peak corresponding to tetramer. Interestingly, there is an appearance of oligomeric species of higher order in the sample incubated with Baicalein. These findings strongly suggest that Baicalein exhibit inhibition activity by sequestration of Tau into oligomers of different nature and taking it off the pathway of aggregation. Although effective against inhibiting fibrillization, Baicalein also worked as a potent molecule for disaggregation of preformed Tau fibrils as well as oligomers (Fig. 7). The ANS fluorescence studies clearly depict the decrease in fluorescence intensity in the initial stages after Baicalein addition, but are found to increase after 12-24 hours at higher concentrations. This behavior of ANS fluorescence can be attributed to the fact that increase in ANS fluorescence depends on binding with hydrophobic residues as well as with cationic amino acids like arginine and lysine ^*72*^. Thus, the Baicalein stabilized oligomer interacts through their cationic residues even though the overall surface hydrophobicity is greatly reduced. The disaggregation potential of Baicalein upon treatment of Tau aggregates signifies that the interaction between these two involves rearrangement of bonds and change in binding energies. If the oligomers formed are stable enough that they do not further interact with other protein species, (either soluble Tau or other oligomers) it can lead to attenuation of further fibrillization of Tau. Altogether, Baicalein plays a dual role of preventing Tau aggregation as well as dissolving the preformed aggregates and can be further considered for the therapeutic screening.

**Figure 7.**
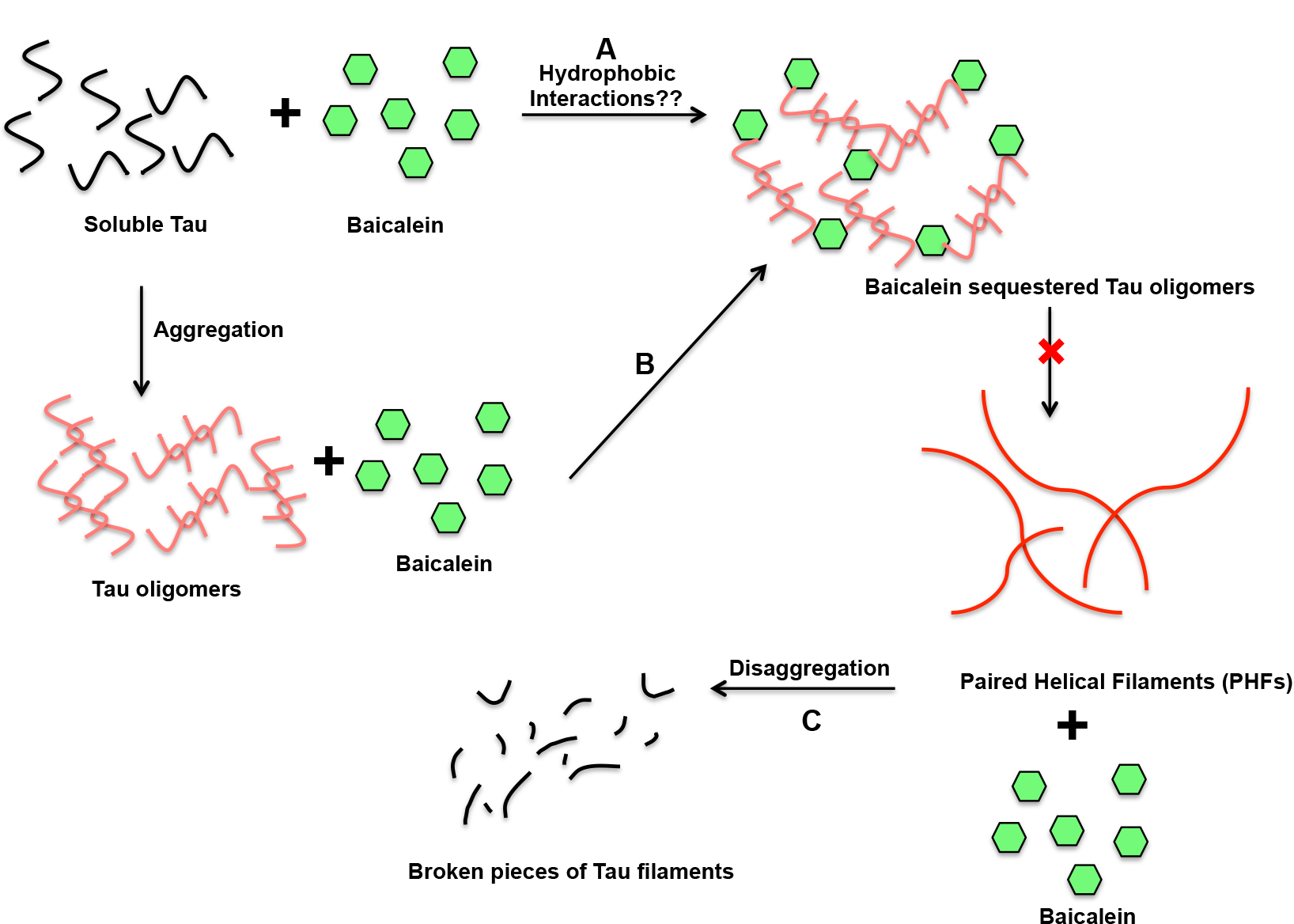
Model for Baicalein-mediated Tau aggregation inhibition and disaggregation. Baicalein may show interaction with various intermittent species of Tau aggregates. A) Baicalein may interact at first step with soluble Tau to induce Tau oligomerization and sequester them based on hydrophobic interactions and prevent their further participation in fibril formation. B) Baicalein might enhance the Tau oligomerization in presence of heparin and further bind these oligomers and inhibit fibril formation. C) Baicalein might also interact with Tau fibrils and lead to their disaggregation leading to formation of small pieces of Tau aggregates.

## Conclusion

The potency of Baicalein in inhibiting the aggregation of proteins like α-synuclein, amyloid beta and amylin has made this molecule important for screening against the other amyloidogenic proteins. Our current investigation demonstrates the ability of Baicalein to inhibit the Tau aggregation by covalent modification. The simulation and docking studies revealed the interaction of 2 adjacent -OH groups of Baicalein with Leucine 266 of the repeat Tau. These adjacent hydroxyl groups might also be involved in the covalent modification of Tau as observed by MALDI-TOF analysis. Along with hydrogen bonding, the hydrophobic interactions were found to play a role in Tau-Baicalein binding. In summary, Baicalein inhibits the Tau aggregation by inducing the off-pathway oligomer formation and preventing the filamentous aggregate formation thus highlighting its potential to be developed as a future therapeutic agent in AD.

## Author contributions

SKS, AB and SC conducted most of the experiments, analyzed the results, and wrote the paper. DB and SKR conducted experiments on Tau-ligand docking and molecular simulation. SC conceived the idea for the project and wrote the paper.

## Acknowledgement

This work was supported in part by grants from the Department of Science and Technology-Science and Engineering Research Board (DST-SERB, Young Investigator grant): SB/YS/LS-355/2013 and DST-SERB/EMR000306, Department of Biotechnology from Neuroscience Task Force (Medical Biotechnology-Human Development & Disease Biology (DBT-HDDB))-BT/PR/15780/MED/30/1629/2015 and in-house CSIR-National Chemical Laboratory grant MLP029526. Shweta Kishor Sonawane acknowledges the fellowship from Department of Biotechnology (DBT), India. Abhishek Ankur Balmik acknowledges the Shyama Prasad Mukherjee fellowship (SPMF) from Council of Scientific Industrial Research (CSIR), India. Tau constructs were kindly gifted by Prof. Roland Brandt from University of Osnabruck, Germany and Prof. Jeff Kuret from Ohio State University College of Medicine, USA. We thank Mr. Yugendra R. Patil, Dr. Santhakumari and Dr. Mahesh J Kulkarni for their fruitful discussion on MALDI-TOF.

## References

[1] Ittner, L. M., and Götz, J. (2011) Amyloid-β and tau—a toxic pas de deux in Alzheimer’s disease, Nature Reviews Neuroscience 12, 67–72.

[2] Hardy, J. (2006) Alzheimer’s disease: the amyloid cascade hypothesis: an update and reappraisal, Journal of Alzheimer’s disease 9, 151–153.

[3] Wang, Y., and Mandelkow, E. (2015) Tau in physiology and pathology, Nature Reviews Neuroscience.

[4] Lovestone, S., Boada, M., Dubois, B., Hüll, M., Rinne, J. O., Huppertz, H.-J., Calero, M., Andrés, M. V., Gómez-Carrillo, B., and León, T. (2015) A phase II trial of tideglusib in Alzheimer’s disease, Journal of Alzheimer’s Disease 45, 75–88.

[5] Morimoto, B. H., Schmechel, D., Hirman, J., Blackwell, A., Keith, J., Gold, M., and Investigators, A.-.-S. (2013) A double-blind, placebo-controlled, ascending-dose, randomized study to evaluate the safety, tolerability and effects on cognition of AL-108 after 12 weeks of intranasal administration in subjects with mild cognitive impairment, Dementia and geriatric cognitive disorders 35, 325–339.

[6] KoSIK, K. S., Joachim, C. L., and Selkoe, D. J. (1986) Microtubule-associated protein tau (tau) is a major antigenic component of paired helical filaments in Alzheimer disease, Proceedings of the National Academy of Sciences 83, 4044–4048.

[7] Kosik, K. S., and McConlogue, L. (1994) Microtubule-associated protein function: Lessons from expression in spodoptera frugiperda cells, Cell motility and the cytoskeleton 28, 195–198.

[8] Ebneth, A., Godemann, R., Stamer, K., Illenberger, S., Trinczek, B., Mandelkow, E.-M., and Mandelkow, E. (1998) Overexpression of tau protein inhibits kinesin-dependent trafficking of vesicles, mitochondria, and endoplasmic reticulum: implications for Alzheimer’s disease, The Journal of cell biology 143, 777–794.

[9] Mandelkow, E.-M., and Mandelkow, E. (2012) Biochemistry and cell biology of tau protein in neurofibrillary degeneration, Cold Spring Harbor perspectives in medicine 2, a006247.

[10] Iqbal, K., Liu, F., Gong, C.-X., Alonso, A. d. C., and Grundke-Iqbal, I. (2009) Mechanisms of tau-induced neurodegeneration, Acta neuropathologica 118, 53–69.

[11] Mandelkow, E.-M., and Mandelkow, E. (1998) Tau in Alzheimer’s disease, Trends in cell biology 8, 425–427.

[12] Cowan, C. M., Quraishe, S., and Mudher, A. (2012) What is the pathological significance of tau oligomers?, Biochemical Society Transactions 40, 693–697.

[13] Bulic, B., Pickhardt, M., Schmidt, B., Mandelkow, E. M., Waldmann, H., and Mandelkow, E. (2009) Development of tau aggregation inhibitors for Alzheimer’s disease, Angewandte Chemie International Edition 48, 1740–1752.

[14] Bulic, B., Pickhardt, M., Mandelkow, E.-M., and Mandelkow, E. (2010) Tau protein and tau aggregation inhibitors, Neuropharmacology 59, 276–289.

[15] Lawatscheck, C., Pickhardt, M., Wieczorek, S., Grafmüller, A., Mandelkow, E., and Börner, H. G. (2016) Generalizing the Concept of Specific Compound Formulation Additives towards Non-Fluorescent Drugs: A Solubilization Study on Potential Anti-Alzheimer-Active Small-Molecule Compounds, Angewandte Chemie International Edition 55, 8752–8756.

[16] Holtzman, D. M., Carrillo, M. C., Hendrix, J. A., Bain, L. J., Catafau, A. M., Gault, L. M., Goedert, M., Mandelkow, E., Mandelkow, E.-M., and Miller, D. S. (2016) Tau: From research to clinical development, Alzheimer’s & Dementia 12, 1033–1039.

[17] Lee, V. M., Brunden, K. R., Hutton, M., and Trojanowski, J. Q. (2011) Developing therapeutic approaches to tau, selected kinases, and related neuronal protein targets, Cold Spring Harbor perspectives in medicine 1, a006437.

[18] Crowe, A., James, M. J., Lee, V. M.-Y., Smith, A. B., Trojanowski, J. Q., Ballatore, C., and Brunden, K. R. (2013) Aminothienopyridazines and methylene blue affect Tau fibrillization via cysteine oxidation, Journal of Biological Chemistry 288, 11024–11037.

[19] Wischik, C., Edwards, P., Lai, R., Roth, M., and Harrington, C. (1996) Selective inhibition of Alzheimer disease-like tau aggregation by phenothiazines, Proceedings of the National Academy of Sciences 93, 11213–11218.

[20] Schirmer, R. H., Adler, H., Pickhardt, M., and Mandelkow, E. (2011) Lest we forget you—methylene blue…, Neurobiology of aging 32, 2325. e2327–2325. e2316.

[21] O’Leary, J. C., Li, Q., Marinec, P., Blair, L. J., Congdon, E. E., Johnson, A. G., Jinwal, U. K., Koren, J., Jones, J. R., and Kraft, C. (2010) Phenothiazine-mediated rescue of cognition in tau transgenic mice requires neuroprotection and reduced soluble tau burden, Molecular neurodegeneration 5, 1.

[22] Wischik, C. M., Harrington, C. R., and Storey, J. M. (2014) Tau-aggregation inhibitor therapy for Alzheimer’s disease, Biochemical pharmacology 88, 529–539.

[23] Akoury, E., Pickhardt, M., Gajda, M., Biernat, J., Mandelkow, E., and Zweckstetter, M. (2013) Mechanistic Basis of Phenothiazine-Driven Inhibition of Tau Aggregation, Angewandte Chemie International Edition 52, 3511–3515.

[24] Ballatore, C., Brunden, K. R., Huryn, D. M., Trojanowski, J. Q., Lee, V. M.-Y., and Smith III, A. B. (2012) Microtubule stabilizing agents as potential treatment for Alzheimer’s disease and related neurodegenerative tauopathies, J. Med. Chem 55, 8979–8996.

[25] Zhang, B., Carroll, J., Trojanowski, J. Q., Yao, Y., Iba, M., Potuzak, J. S., Hogan, A.-M. L., Xie, S. X., Ballatore, C., and Smith, A. B. (2012) The microtubule-stabilizing agent, epothilone D, reduces axonal dysfunction, neurotoxicity, cognitive deficits, and Alzheimer-like pathology in an interventional study with aged tau transgenic mice, Journal of Neuroscience 32, 3601–3611.

[26] Brunden, K. R., Zhang, B., Carroll, J., Yao, Y., Potuzak, J. S., Hogan, A.-M. L., Iba, M., James, M. J., Xie, S. X., and Ballatore, C. (2010) Epothilone D improves microtubule density, axonal integrity, and cognition in a transgenic mouse model of tauopathy, Journal of Neuroscience 30, 13861–13866.

[27] Calcul, L., Zhang, B., Jinwal, U. K., Dickey, C. A., and Baker, B. J. (2012) Natural products as a rich source of tau-targeting drugs for Alzheimer’s disease, Future medicinal chemistry 4, 1751–1761.

[28] Paranjape, S. R., Riley, A. P., Somoza, A. D., Oakley, C. E., Wang, C. C., Prisinzano, T. E., Oakley, B. R., and Gamblin, T. C. (2015) Azaphilones inhibit tau aggregation and dissolve tau aggregates in vitro, ACS chemical neuroscience 6, 751–760.

[29] Paranjape, S. R., Chiang, Y.-M., Sanchez, J. F., Entwistle, R., Wang, C. C., Oakley, B. R., and Gamblin, T. C. (2014) Inhibition of Tau aggregation by three Aspergillus nidulans secondary metabolites: 2, ω-dihydroxyemodin, asperthecin, and asperbenzaldehyde, Planta medica 80, 77–85.

[30] Li, W., Sperry, J. B., Crowe, A., Trojanowski, J. Q., Smith III, A. B., and Lee, V. M. Y. (2009) Inhibition of tau fibrillization by oleocanthal via reaction with the amino groups of tau, Journal of neurochemistry 110, 1339–1351.

[31] Bieschke, J., Russ, J., Friedrich, R. P., Ehrnhoefer, D. E., Wobst, H., Neugebauer, K., and Wanker, E. E. (2010) EGCG remodels mature α-synuclein and amyloid-β fibrils and reduces cellular toxicity, Proceedings of the National Academy of Sciences 107, 7710–7715.

[32] Wobst, H. J., Sharma, A., Diamond, M. I., Wanker, E. E., and Bieschke, J. (2015) The green tea polyphenol (−)-epigallocatechin gallate prevents the aggregation of tau protein into toxic oligomers at substoichiometric ratios, FEBS letters 589, 77–83.

[33] Vauzour, D., Vafeiadou, K., Rodriguez-Mateos, A., Rendeiro, C., and Spencer, J. P. (2008) The neuroprotective potential of flavonoids: a multiplicity of effects, Genes & nutrition 3, 115–126.

[34] Zhu, M., Rajamani, S., Kaylor, J., Han, S., Zhou, F., and Fink, A. L. (2004) The flavonoid baicalein inhibits fibrillation of α-synuclein and disaggregates existing fibrils, Journal of Biological Chemistry 279, 26846–26857.

[35] Zhang, S. Q., Obregon, D., Ehrhart, J., Deng, J., Tian, J., Hou, H., Giunta, B., Sawmiller, D., and Tan, J. (2013) Baicalein reduces β-amyloid and promotes nonamyloidogenic amyloid precursor protein processing in an Alzheimer’s disease transgenic mouse model, Journal of neuroscience research 91, 1239–1246.

[36] Huy, P. D. Q., Vuong, Q. V., La Penna, G., Faller, P., and Li, M. S. (2016) Impact of Cu (II) binding on Structures and Dynamics of Aβ42 Monomer and Dimer: Molecular Dynamics Study, ACS Chemical Neuroscience 7, 1348–1363.

[37] Barghorn, S., Biernat, J., and Mandelkow, E. (2005) Purification of recombinant tau protein and preparation of Alzheimer-paired helical filaments in vitro, Amyloid Proteins: Methods and Protocols, 35–51.

[38] Altschul, S. F., Gish, W., Miller, W., Myers, E. W., and Lipman, D. J. (1990) Basic local alignment search tool, Journal of molecular biology 215, 403–410.

[39] Webb, B., and Sali, A. (2014) Protein structure modeling with MODELLER, In Protein Structure Prediction, pp 1–15, Springer.

[40] Laskowski, R. A., MacArthur, M. W., Moss, D. S., and Thornton, J. M. (1993) PROCHECK: a program to check the stereochemical quality of protein structures, Journal of applied crystallography 26, 283–291.

[41] Lovell, S. C., Davis, I. W., Arendall III, W. B., De Bakker, P. I., Word, J. M., Prisant, M. G., Richardson, J. S., and Richardson, D. C. (2003) Structure validation by Cα geometry: ϕ, Ψ and Cβ deviation, Proteins: Structure, Function, and Bioinformatics 50, 437–450.

[42] Colovos, C., and Yeates, T. O. (1993) Verification of protein structures: patterns of nonbonded atomic interactions, Protein science 2, 1511–1519.

[43] Wiederstein, M., and Sippl, M. J. (2007) ProSA-web: interactive web service for the recognition of errors in three-dimensional structures of proteins, Nucleic acids research 35, W407–W410.

[44] Sastry, G. M., Adzhigirey, M., Day, T., Annabhimoju, R., and Sherman, W. (2013) Protein and ligand preparation: parameters, protocols, and influence on virtual screening enrichments, Journal of computer-aided molecular design 27, 221–234.

[45] Friesner, R. A., Banks, J. L., Murphy, R. B., Halgren, T. A., Klicic, J. J., Mainz, D. T., Repasky, M. P., Knoll, E. H., Shelley, M., and Perry, J. K. (2004) Glide: a new approach for rapid, accurate docking and scoring. 1. Method and assessment of docking accuracy, Journal of medicinal chemistry 47, 1739–1749.

[46] Halgren, T. A. (2009) Identifying and characterizing binding sites and assessing druggability, Journal of chemical information and modeling 49, 377–389.

[47] Bowers, K. J., Chow, D. E., Xu, H., Dror, R. O., Eastwood, M. P., Gregersen, B. A., Klepeis, J. L., Kolossvary, I., Moraes, M. A., and Sacerdoti, F. D. (2006) Scalable algorithms for molecular dynamics simulations on commodity clusters, In SC’06: Proceedings of the 2006 ACM/IEEE Conference on Supercomputing, pp 43–43, IEEE.

[48] Jeganathan, S., von Bergen, M., Brutlach, H., Steinhoff, H.-J., and Mandelkow, E. (2006) Global hairpin folding of tau in solution, Biochemistry 45, 2283–2293.

[49] Chenprakhon, P., Sucharitakul, J., Panijpan, B., and Chaiyen, P. (2010) Measuring Binding Affinity of Protein-Ligand Interaction Using Spectrophotometry: Binding of Neutral Red to Riboflavin-Binding Protein, Journal of chemical education 87, 829–831.

[50] Hu, Y.-J., Liu, Y., Wang, J.-B., Xiao, X.-H., and Qu, S.-S. (2004) Study of the interaction between monoammonium glycyrrhizinate and bovine serum albumin, Journal of Pharmaceutical and Biomedical Analysis 36, 915–919.

[51] Zsila, F., Bikádi, Z., and Simonyi, M. (2003) Probing the binding of the flavonoid, quercetin to human serum albumin by circular dichroism, electronic absorption spectroscopy and molecular modelling methods, Biochemical pharmacology 65, 447–456.

[52] Daebel, V., Chinnathambi, S., Biernat, J., Schwalbe, M., Habenstein, B., Loquet, A., Akoury, E., Tepper, K., Müller, H., and Baldus, M. (2012) β-Sheet core of tau paired helical filaments revealed by solid-state NMR, Journal of the American Chemical Society 134, 13982–13989.

[53] Mukrasch, M. D., Biernat, J., von Bergen, M., Griesinger, C., Mandelkow, E., and Zweckstetter, M. (2005) Sites of tau important for aggregation populate {beta}-structure and bind to microtubules and polyanions, The Journal of biological chemistry 280, 24978–24986.

[54] Von Bergen, M., Barghorn, S., Biernat, J., Mandelkow, E.-M., and Mandelkow, E. (2005) Tau aggregation is driven by a transition from random coil to beta sheet structure, Biochimica et Biophysica Acta (BBA)-Molecular Basis of Disease 1739, 158–166.

[55] Friedhoff, P., Von Bergen, M., Mandelkow, E.-M., Davies, P., and Mandelkow, E. (1998) A nucleated assembly mechanism of Alzheimer paired helical filaments, Proceedings of the National Academy of Sciences 95, 15712–15717.

[56] Goedert, M., Jakes, R., Spillantini, M., Hasegawa, M., Smith, M., and Crowther, R. (1996) Assembly of microtubule-associated protein tau into Alzheimer-like filaments induced by sulphated glycosaminoglycans.

[57] Pickhardt, M., Gazova, Z., von Bergen, M., Khlistunova, I., Wang, Y., Hascher, A., Mandelkow, E.-M., Biernat, J., and Mandelkow, E. (2005) Anthraquinones inhibit tau aggregation and dissolve Alzheimer’s paired helical filaments in vitro and in cells, Journal of Biological Chemistry 280, 3628–3635.

[58] Ramachandran, G., and Udgaonkar, J. B. (2011) Understanding the kinetic roles of the inducer heparin and of rod-like protofibrils during amyloid fibril formation by Tau protein, Journal of Biological Chemistry 286, 38948–38959.

[59] Kumar, S., Tepper, K., Kaniyappan, S., Biernat, J., Wegmann, S., Mandelkow, E.-M., Müller, D. J., and Mandelkow, E. (2014) Stages and conformations of the Tau repeat domain during aggregation and its effect on neuronal toxicity, Journal of Biological Chemistry 289, 20318–20332.

[60] Crowe, A., Ballatore, C., Hyde, E., Trojanowski, J. Q., and Lee, V. M.-Y. (2007) High throughput screening for small molecule inhibitors of heparin-induced tau fibril formation, Biochemical and biophysical research communications 358, 1–6.

[61] Brunden, K. R., Trojanowski, J. Q., and Lee, V. M.-Y. (2009) Advances in tau-focused drug discovery for Alzheimer’s disease and related tauopathies, Nature reviews Drug discovery 8, 783–793.

[62] Bulic, B., Pickhardt, M., and Mandelkow, E. (2013) Progress and developments in tau aggregation inhibitors for Alzheimer disease, J. Med. Chem 56, 4135–4155.

[63] Congdon, E. E., Kim, S., Bonchak, J., Songrug, T., Matzavinos, A., and Kuret, J. (2008) Nucleation-dependent tau filament formation the importance of dimerization and an estimation of elementary rate constants, Journal of Biological Chemistry 283, 13806–13816.

[64] Flach, K., Hilbrich, I., Schiffmann, A., Gärtner, U., Krüger, M., Leonhardt, M., Waschipky, H., Wick, L., Arendt, T., and Holzer, M. (2012) Tau oligomers impair artificial membrane integrity and cellular viability, Journal of Biological Chemistry 287, 43223–43233.

[65] Guzmán-Martinez, L., Farías, G. A., and Maccioni, R. B. (2014) Tau oligomers as potential targets for Alzheimer’s diagnosis and novel drugs, Tau oligomers 3, 35.

[66] Chua, S. W., Cornejo, A., van Eersel, J., Stevens, C. H., Vaca, I., Cueto, M., Kassiou, M., Gladbach, A., Macmillan, A., and Lewis, L. (2017) The Polyphenol Altenusin Inhibits in Vitro Fibrillization of Tau and Reduces Induced Tau Pathology in Primary Neurons, ACS chemical neuroscience 8, 743–751.

[67] Margittai, M., and Langen, R. (2004) Template-assisted filament growth by parallel stacking of tau, Proceedings of the National Academy of Sciences of the United States of America 101, 10278–10283.

[68] Semisotnov, G., Rodionova, N., Razgulyaev, O., Uversky, V., Gripas, A., and Gilmanshin, R. (1991) Study of the “molten globule” intermediate state in protein folding by a hydrophobic fluorescent probe, Biopolymers 31, 119–128.

[69] Taniguchi, S., Suzuki, N., Masuda, M., Hisanaga, S.-i., Iwatsubo, T., Goedert, M., and Hasegawa, M. (2005) Inhibition of heparin-induced tau filament formation by phenothiazines, polyphenols, and porphyrins, Journal of Biological Chemistry 280, 7614–7623.

[70] Monti, M. C., Margarucci, L., Riccio, R., and Casapullo, A. (2012) Modulation of tau protein fibrillization by oleocanthal, Journal of natural products 75, 1584–1588.

[71] Cisek, K., L Cooper, G., J Huseby, C., and Kuret, J. (2014) Structure and mechanism of action of tau aggregation inhibitors, Current Alzheimer Research 11, 918–927.

[72] Gasymov, O. K., and Glasgow, B. J. (2007) ANS fluorescence: Potential to augment the identification of the external binding sites of proteins, Biochimica et Biophysica Acta (BBA)-Proteins and Proteomics 1774, 403–411.

