## Supplementary material for "Baicalein suppresses Tau fibrillization by sequestering oligomers": SI

#### Supporting Figures

### Supplementary Figure 1

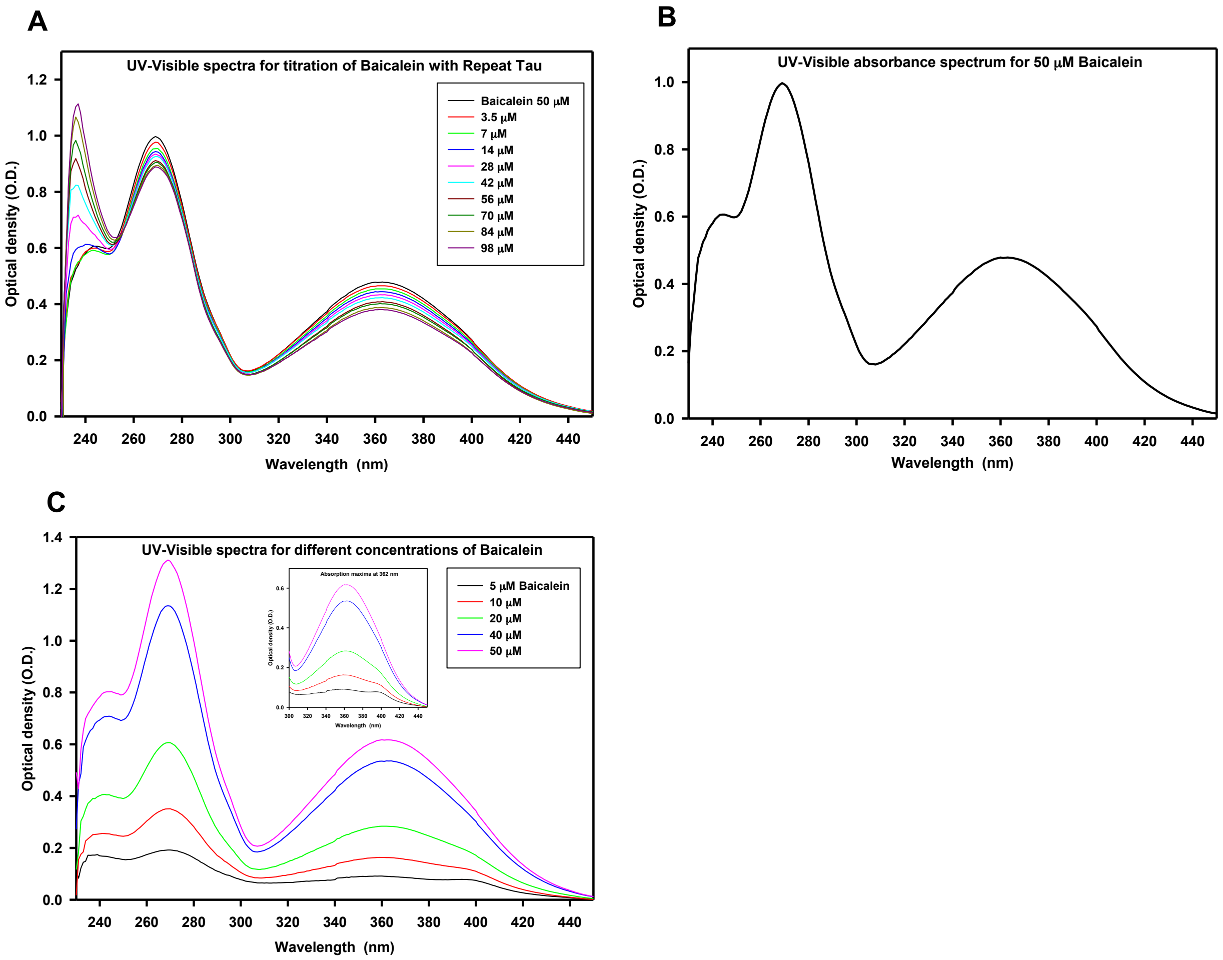


**Figure S1. UV –Visible absorption spectra of Baicalein.** A) The UV-VIS absorbance spectra for Baicalein obtained by titrating 50 µM of Baicalein with different concentrations of repeat Tau (3.5, 7, 14, 28, 42, 56, 70, 84, 98 µM). B) UV-VIS absorption spectrum for 50 µM Baicalein in 20 mM BES buffer, pH 7.4 showing maxima at 270 nm and 362 nm. C) Absorption spectra for different concentrations of Baicalein.

#### Supplementary Figure 2

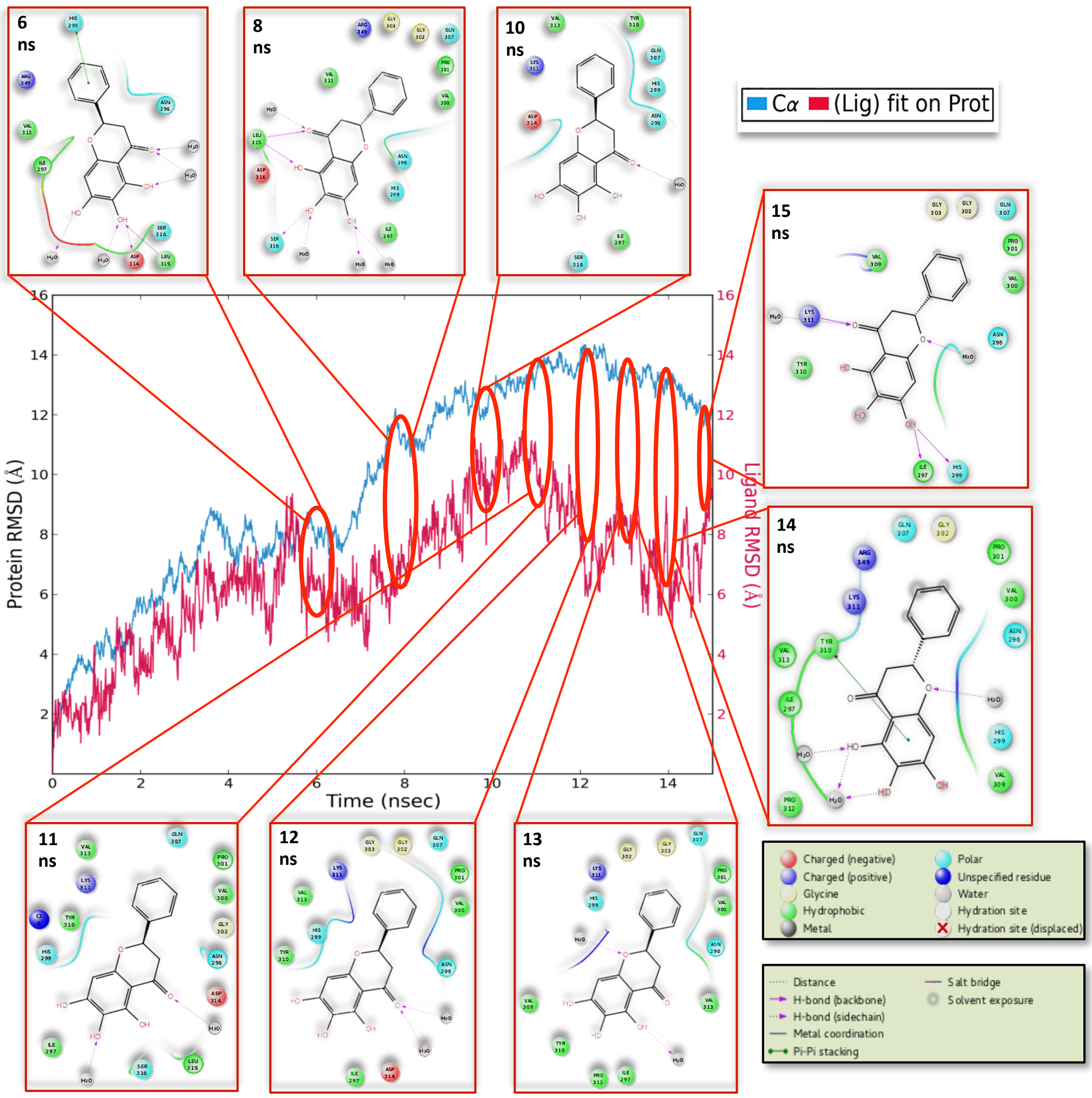


**Figure S2. RMSD plot of Tau-Baicalein interaction.** The above plot shows the RMSD evolution of the protein, Tau (left *Y*-axis) and a comparative view of the RMSD evolution of the ligand, Baicalein (right *Y*-axis) through the similar time-scale, which indicates how stable the Baicalein is with respect to the binding pocket in Tau. The interactions are shown in detail, for different nanoseconds for comparative relevance. The diagram indicates a dynamic pocket of interaction of Tau, which is to be expected since Tau is a highly flexible soluble protein.

#### Supplementary Figure 3

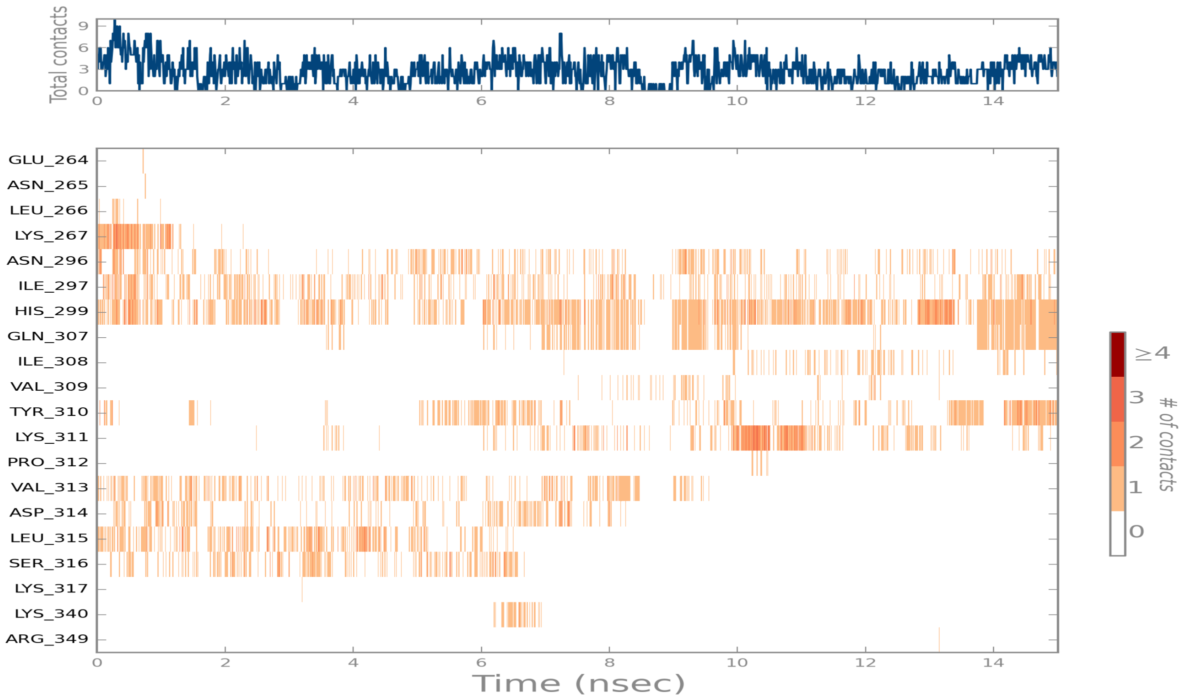


**Figure S3. A timeline representation of the interactions and contacts (H-bonds, Hydrophobic, Ionic, Water bridges) over the entire timeline of the interaction**. The top panel shows the total number of specific contacts the Tau makes with the Baicalein over the course of the trajectory. The bottom panel shows which residues interact with the Baicalein in each trajectory frame. Some residues of Tau make more than one specific contact with the Baicalein, which is represented by a darker shade of orange, according to the scale to the right of the plot.

#### Supplementary Figure 4

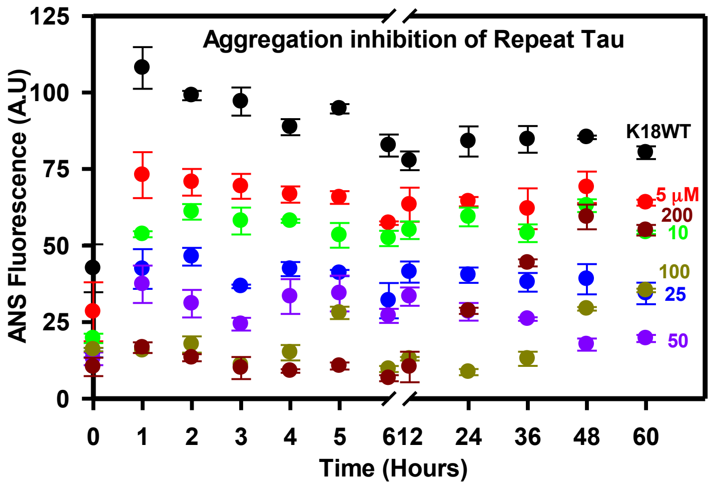


**Figure S4. The ANS fluorescence analysis of repeat Tau aggregation inhibition by Baicalein.** The ANS analysis of Baicalein treated Tau from 5-200 μM shows initial decrease in the fluorescence followed by increase at the later time points suggesting increase in Tau hydrophobicity.

#### Supplementary Figure 5

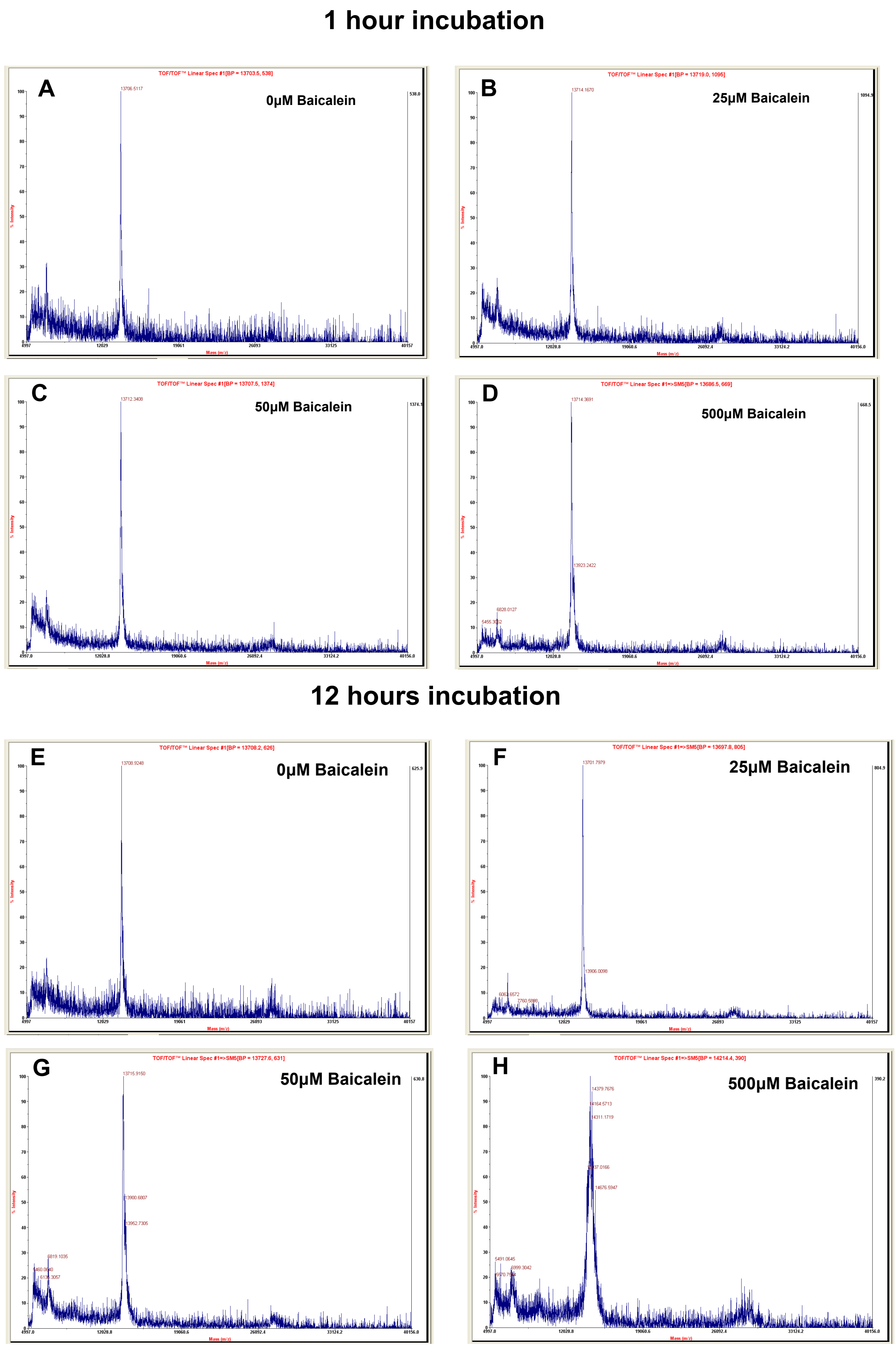


**Figure S5.** MALDI analysis of Baicalein and repeat Tau. A, B, C, D) The MALDI analysis of repeat Tau with different concentrations of Baicalein incubated for 1 hour. E, F, G, H) The MALDI analysis of repeat Tau with different concentrations of Baicalein incubated for 12 hours.

### Supplementary Figure 6

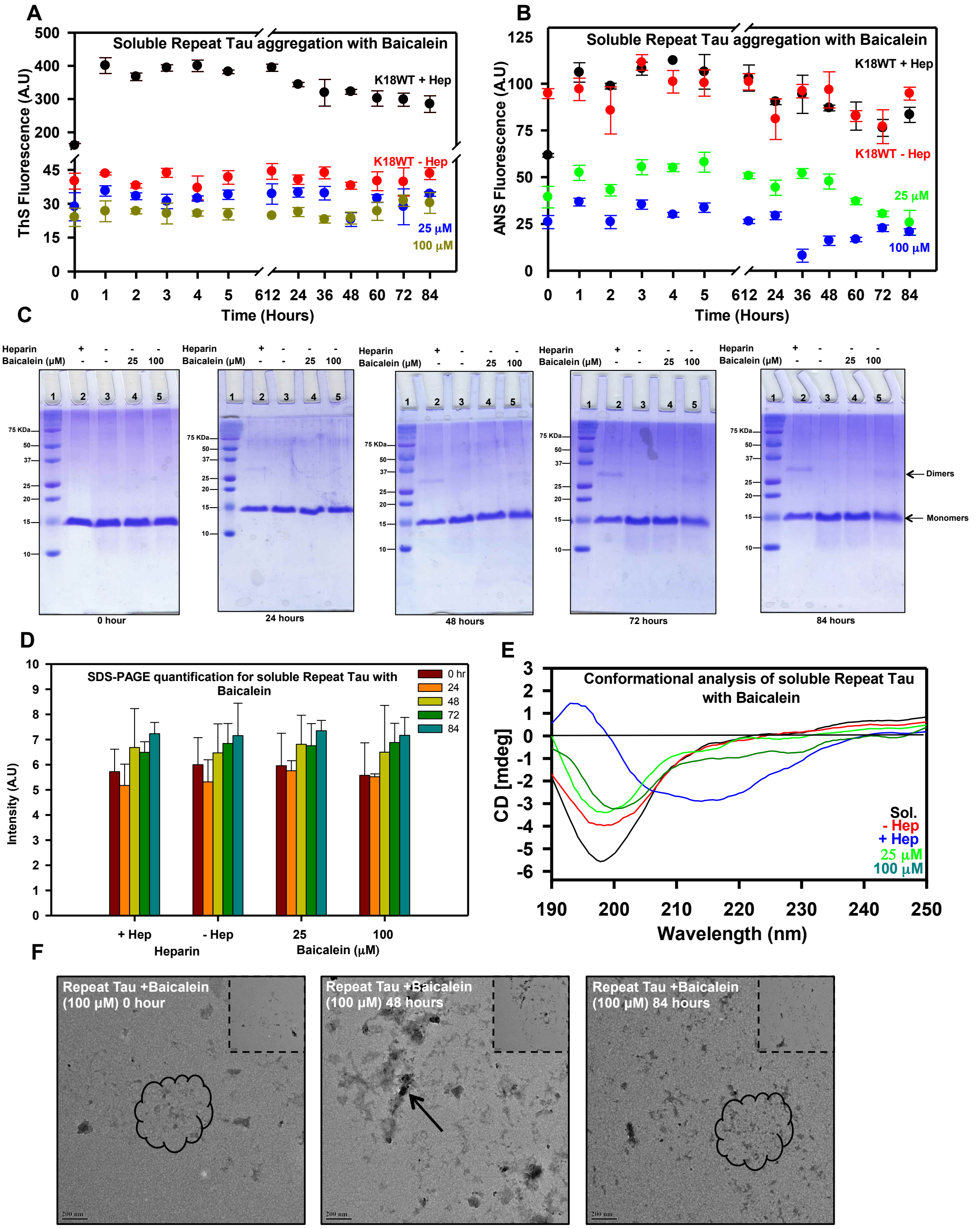


**Figure S6. Effect of Baicalein on repeat Tau aggregation in absence of heparin.** A) ThS fluorescence show increase in intensity in the heparin treated positive control whereas no increase in fluorescence was observed in the negative control as well as Baicalein treated samples. B) Baicalein does not increase the hydrophobicity as revealed by ANS fluorescence. C) SDS-PAGE analysis shows the presence of dimer and tetramers in the positive control from 24 hours onwards (lane 2). A faint band of dimer and a tetramer is observed in 100 µM Baicalein treated sample from 48 hours onwards but has a lesser intensity as compared to positive control. D) Quantification for SDS-PAGE for soluble repeat Tau with Baicalein. CD analysis for the effect of Baicalein on repeat Tau in absence of heparin. E) The CD spectra shows shift towards beta sheet whereas the Baicalein treated samples shows the spectra of a typically random coiled protein. F) Amorphous aggregates of repeat Tau formed at different time points in presence of Baicalein as observed by electron microscopy.
